## Supplementary material for "AMODO-EO: A Practical Framework for Adaptive Objective Discovery in Multi-Objective Drug Optimization": Source code of the algorithm: CHANGELOG.md

# Changelog

## [1.0.0] - 2025-08-16

### Added
- Initial release of AMODO-EO algorithm
- Core adaptive multi-objective optimization framework
- Statistical validation and benchmarking tools
- Experimental validation on ChEMBL datasets
- Complete manuscript and documentation

### Features
- Adaptive objective discovery during optimization
- Statistical independence and variance validation
- Chemical interpretability filtering
- Performance benchmarking against NSGA-II
- Comprehensive ablation studies
