## Supplementary material for "AMODO-EO: A Practical Framework for Adaptive Objective Discovery in Multi-Objective Drug Optimization": Source code of the algorithm: CONTRIBUTING.md

# Contributing to AMODO-EO

We welcome contributions to the AMODO-EO project! Please follow these guidelines:

## How to Contribute

1. Fork the repository
2. Create a feature branch (`git checkout -b feature/amazing-feature`)
3. Commit your changes (`git commit -m 'Add amazing feature'`)
4. Push to the branch (`git push origin feature/amazing-feature`)
5. Open a Pull Request

## Development Guidelines

- Follow PEP 8 style guidelines for Python code
- Include docstrings for all functions and classes
- Add unit tests for new functionality
- Update documentation as needed

## Issues

Please use the GitHub issue tracker to report bugs or request features.

## Code of Conduct

This project follows a standard code of conduct. Please be respectful and constructive in all interactions.
