## Supplementary material for "AMODO-EO: A Practical Framework for Adaptive Objective Discovery in Multi-Objective Drug Optimization": Source code of the algorithm: README.md

# AMODO-EO: Adaptive Multi-Objective Drug Optimization with Emergent Objectives

This repository contains the implementation and manuscript for "AMODO-EO: A Practical Framework for Adaptive Objective Discovery in Multi-Objective Drug Optimization" by Shetty Nihal Naveen, B. Karthik Udupa, and Ashil A. Shetty.

## 📄 Manuscript

The main manuscript is available in LaTeX format:
- `manuscript/amodo_heo_comprehensive_research_paper.tex` - Main manuscript file
- `manuscript/references.bib` - Bibliography file

## 🔬 Algorithm Implementation

The core AMODO-EO algorithm implementation:
- `src/publication_ready_amodo_heo.py` - Main algorithm implementation
- `src/statistical_benchmarking.py` - Benchmarking and statistical validation

## 🧪 Experiments

Experimental validation code:
- `experiments/experimental_validation.py` - Experimental validation framework

## 📊 Results

Key experimental results:
- `results/final_objectives.json` - Final discovered objectives and performance metrics

## 🚀 Getting Started

### Prerequisites
```bash
pip install -r requirements.txt
```

### Running the Algorithm
```python
from src.publication_ready_amodo_heo import AMODEO

# Initialize and run AMODO-EO
optimizer = AMODEO(population_size=100, generations=500)
results = optimizer.optimize(dataset)
```

## 📜 Citation

If you use this work, please cite:
```bibtex
@article{shetty2025amodo,
  title={AMODO-EO: A Practical Framework for Adaptive Objective Discovery in Multi-Objective Drug Optimization},
  author={Shetty, Nihal Naveen and Udupa, B. Karthik and Shetty, Ashil A.},
  journal={arXiv preprint},
  year={2025}
}
```

## 📧 Contact

- Shetty Nihal Naveen:
- B. Karthik Udupa:  
- Ashil A. Shetty:

## 🏫 Affiliation

Department of Information Science & Engineering  
Sahyadri College of Engineering and Management  
Mangaluru, Karnataka, India

## 📄 License

This project is available under MIT License.
