## Supplementary material for "AMODO-EO: A Practical Framework for Adaptive Objective Discovery in Multi-Objective Drug Optimization": Source code of the algorithm: REAL_DATA_PLAN.md

# REAL DATA ENHANCEMENT PLAN
# For future journal submission after arXiv

## Quick Real Data Options (1-2 weeks):

### 1. Public ChEMBL Data (Free & Easy)
- Download from: https://www.ebi.ac.uk/chembl/
- 2M+ bioactivity measurements
- Already preprocessed
- Standard in drug discovery research

### 2. PubChem Data (Free & Massive)
- 100M+ chemical compounds
- Bioactivity data available
- Easy API access

### 3. FDA Orange Book (Regulatory Data)
- Approved drug data
- Bioequivalence information
- Publicly available

## Implementation Plan:

### Week 1: Data Collection
- Download ChEMBL bioactivity data (IC50, Ki values)
- Extract molecular descriptors using RDKit
- Create 3-4 real objective functions

### Week 2: Validation
- Run AMODO-EO on real data
- Compare with synthetic results
- Update manuscript with real validation

## Advantages of This Approach:
1. **ArXiv first** - Establishes priority and gets feedback
2. **Real data later** - Strengthens for journal submission
3. **Both validations** - Shows robustness across data types
