## Supplementary material for "AMODO-EO: A Practical Framework for Adaptive Objective Discovery in Multi-Objective Drug Optimization": Source code of the algorithm: setup.py

```
from setuptools import setup, find_packages

with open("README.md", "r", encoding="utf-8") as fh:
    long_description = fh.read()

with open("requirements.txt", "r", encoding="utf-8") as fh:
    requirements = [line.strip() for line in fh if line.strip() and not line.startswith("#")]

setup(
    name="amodo-eo",
    version="1.0.0",
    author="Shetty Nihal Naveen, B. Karthik Udupa, Ashil A. Shetty",
    author_email="",
    description="AMODO-EO: A Practical Framework for Adaptive Objective Discovery in Multi-Objective Drug Optimization",
    long_description=long_description,
    long_description_content_type="text/markdown",
    url="https://github.com/nihsh2004/AMODO-EO",
    packages=find_packages(),
    classifiers=[
        "Development Status :: 4 - Beta",
        "Intended Audience :: Science/Research",
        "License :: OSI Approved :: MIT License",
        "Operating System :: OS Independent",
        "Programming Language :: Python :: 3",
        "Programming Language :: Python :: 3.8",
        "Programming Language :: Python :: 3.9",
        "Programming Language :: Python :: 3.10",
        "Topic :: Scientific/Engineering :: Artificial Intelligence",
        "Topic :: Scientific/Engineering :: Chemistry",
    ],
    python_requires=">=3.8",
    install_requires=requirements,
    keywords="multi-objective optimization, drug discovery, adaptive algorithms, evolutionary computation",
)
```
