## Supplementary material for "AMODO-EO: A Practical Framework for Adaptive Objective Discovery in Multi-Objective Drug Optimization": Source code of the algorithm: amodo_heo_comprehensive_research_paper.tex

% AMODO-EO: Adaptive Multi-Objective Drug Optimization with Emergent Objectives
% Authors: Shetty Nihal Naveen, B. Karthik Udupa, Ashil A. Shetty
% Institution: Sahyadri College of Engineering and Management, India
% arXiv submission: August 2025

\documentclass[11pt,a4paper]{article}
\usepackage[utf8]{inputenc}
\usepackage[T1]{fontenc}
\usepackage{amsmath,amssymb,amsfonts}
\usepackage{algorithm}
\usepackage{algorithmic}
\usepackage{graphicx}
\usepackage{booktabs}
\usepackage{array}
\usepackage{multirow}
\usepackage{float}
\usepackage{hyperref}
\usepackage{natbib}
\usepackage{geometry}
\usepackage{xcolor}
\usepackage{caption}
\usepackage{subcaption}
\usepackage{setspace}

% Set proper margins for arXiv
\geometry{
    margin=2cm,
    top=2.5cm,
    bottom=2.5cm
}

% Set line spacing (single spacing preferred for arXiv)
\singlespacing

% Custom commands for definitions
\newtheorem{definition}{Definition}
\newtheorem{theorem}{Theorem}

% Title and author information
\title{AMODO-EO: A Practical Framework for Adaptive Objective Discovery in Multi-Objective Drug Optimization}

\author{
Shetty Nihal Naveen$^{1}$, B. Karthik Udupa$^{1}$, Ashil A. Shetty$^{1}$\\
\\
$^{1}$Department of Information Science \& Engineering,\\
Sahyadri College of Engineering and Management,\\
Mangaluru, Karnataka, India\\
\\
\texttt{}\\
\texttt{}\\
\texttt{}
}

\date{August 2025}

\begin{document}

\maketitle

\begin{abstract}
Multi-objective optimization in drug discovery faces a fundamental limitation: the assumption that all relevant optimization criteria can be predefined. Traditional approaches using fixed objective sets often miss critical molecular trade-offs that emerge during the optimization process. We present AMODO-EO (Adaptive Multi-Objective Drug Optimization with Emergent Objectives), a novel framework that systematically discovers, validates, and integrates new optimization criteria during the search process by analyzing chemical data patterns. Building upon established automated feature engineering principles, AMODO-EO extends these techniques to dynamic multi-objective optimization contexts in drug discovery. The framework automatically generates candidate objectives from molecular descriptor relationships, validates them through rigorous statistical independence and variance criteria, and integrates them into the optimization process using adaptive weighting mechanisms. Comprehensive experiments on ChEMBL datasets demonstrate consistent identification of chemically meaningful objectives such as the hydrogen-bond acceptor/rotatable-bond ratio, which captures previously unspecified trade-offs between molecular polarity and flexibility. Statistical validation confirms that discovered objectives are independent of predefined criteria while providing significant discriminative power. Performance analysis shows that AMODO-EO successfully extends multi-objective formulations while maintaining competitive optimization performance on original objectives, with only 2-3\% performance cost for the discovery capability. The framework opens new possibilities for adaptive optimization in chemical design by automatically revealing hidden molecular relationships that provide actionable insights for drug designers. Our approach represents a significant advancement in computational drug discovery by transforming multi-objective optimization from a purely exploitative process into one that systematically explores the space of potential optimization criteria.
\end{abstract}

\textbf{Keywords:} Multi-objective optimization, Drug discovery, Adaptive algorithms, Emergent objectives, Objective discovery, Mathematical programming

\section{Introduction}

Multi-objective optimization (MOO) in drug discovery must balance conflicting goals (e.g., potency vs. safety) over a vast chemical search space \citep{lipinski2001experimental,veber2002molecular}. Traditional MOO methods (e.g., NSGA-II \citep{deb2002fast}, MOEA/D \citep{zhang2007moead}) assume a fixed objective set defined \emph{a priori}. However, real-world molecular design often involves hidden trade-offs among chemical descriptors that are not captured by the initial objectives \citep{bohacek1996art,reymond2015chemical}. Recent dynamic MOO approaches can adapt to changing objectives, but typically assume those changes are externally imposed \citep{jin2005evolutionary}. 

Despite significant advances in multi-objective optimization for drug discovery \citep{brown2019guacamol,polykovskiy2020molecular}, a fundamental limitation persists: the assumption that all relevant optimization criteria can be identified and defined before the search begins. This constraint forces researchers to either rely on incomplete objective sets that may miss critical molecular trade-offs, or attempt to anticipate all possible design criteria, which is often impractical given the complexity of chemical space \citep{kirkpatrick1994exploring,fink2007virtual}.

\subsection{Related Work and Positioning}

Our approach builds upon established principles from symbolic regression (SR) and automated feature engineering (AFE), extending these mature techniques to address the distinct challenge of objective discovery during active multi-objective optimization. We acknowledge the foundational work in these areas and position AMODO-EO as a novel integration rather than a paradigm-shifting invention.

\textbf{Symbolic Regression and Automated Discovery:} SR methods like Eureqa \citep{schmidt2009distilling} and AI Feynman \citep{udrescu2020ai} have been cornerstones of automated scientific discovery for over a decade, systematically exploring mathematical expressions to discover relationships in data. These approaches successfully identify physical laws and mathematical relationships across diverse domains by searching vast spaces of functional forms to uncover intrinsic relationships. Traditional SR focuses on finding single expressions that best fit target variables for prediction tasks, operating on static datasets to build predictive models.

\textbf{Automated Feature Engineering:} AFE techniques, well-established in machine learning \citep{khurana2018feature}, automatically generate large numbers of candidate features through mathematical operations on primary descriptors, followed by selection processes to identify the most relevant ones. In cheminformatics, sophisticated AFE approaches using grammar-based genetic programming have been successfully applied to catalyst design \citep{toyao2020machine} and ADMET property prediction \citep{yang2019auto,ertl2009estimation}, exploring rich spaces of potential chemical relationships. These methods represent mature and highly relevant approaches to the fundamental challenge of discovering meaningful mathematical relationships in chemical data \citep{rogers2010extended,morgan1965generation}.

\textbf{Dynamic Multi-Objective Optimization:} Dynamic MOO methods adapt to changing objective landscapes over time \citep{jin2005evolutionary}. Approaches like dynamic NSGA-II and time-varying MOEAs handle objectives that change due to external factors or temporal evolution. However, these methods assume that objective modifications are externally imposed rather than discovered from the optimization data itself.

\textbf{Our Specific Contribution:} AMODO-EO represents a novel application and integration of established AFE/SR principles within the specific context of dynamic multi-objective optimization for drug discovery. Rather than claiming to invent objective discovery, we acknowledge that our core discovery mechanism—generating candidate functions through mathematical operators on descriptor pairs—is functionally equivalent to specific implementations of symbolic regression and automated feature engineering. 

The genuine novelty lies in the integration: (1) performing objective discovery during active Pareto front evolution rather than static feature discovery, (2) maintaining multi-objective optimization performance while dynamically expanding dimensionality, (3) incorporating domain-specific chemical interpretability constraints for drug discovery, (4) using conflict resolution matrices and adaptive weight evolution to manage evolving objective relationships, and (5) balancing exploitation of known objectives with exploration of potential new ones.

This positions our work accurately as a specialized application of mature techniques to an unexplored problem context, rather than a fundamental algorithmic breakthrough. We acknowledge that by building upon existing foundations, we could have explored more sophisticated discovery mechanisms from the AFE/SR literature, which represents a valuable direction for future enhancement.

The literature reveals this gap clearly—while methods exist for adapting to externally imposed objective changes, no systematic framework exists for discovering new, chemically meaningful objectives from the optimization data itself during active search processes.

In this work, we introduce AMODO-EO, a mathematical framework that integrates objective discovery into the MOO process. Rather than limiting the search to a pre-specified objective vector $\mathbf{F}(x) = [f_1(x), \dots, f_k(x)]^T$, AMODO-EO allows a dynamic objective set that grows over time:

\begin{equation}
\mathbf{F}^t(x) = [f_1(x), \dots, f_k(x), e^{(t)}_{1}(x), \dots, e^{(t)}_{m_t}(x)]^T,
\end{equation}

where $e^{(t)}_j(x)$ are emergent objectives discovered up to iteration $t$ (and $m_t$ is their count). Initially $m_0=0$. Our goal is to identify valid new objectives from data and add them to the optimization as it proceeds.

The key idea is to scan the current solution set and historical data for statistically valid and interpretable descriptor relationships that serve as new objectives. We systematically generate candidate functions (e.g., descriptor ratios, products, or differences) and filter them by correlation and variance criteria, only retaining those that are both independent of existing objectives and sufficiently discriminative. Validated candidates are added to the objective set for subsequent generations.

The algorithm then optimizes the expanded objective set using a combination of evolutionary search and adaptive weight learning. We use a (weighted) hypervolume indicator as a proxy objective for learning weight updates: specifically, we maintain a weight vector $\mathbf{w}$ over objectives and update it by (approximate) gradient ascent on the weighted hypervolume of the population, with entropy regularization to encourage weight diversity. Simultaneously, we compute a conflict matrix $C$ from pairwise objective correlations and solve a quadratic program to de-emphasize highly correlated (redundant) objectives. A parameter $\beta$ (set to 0.7) controls the interpolation between these gradient-based and conflict-based updates. Finally, standard genetic operators (crossover, mutation, selection) evolve the population under the current preferences (Algorithm~\ref{alg:amodo}).

Our contributions are: (1) the first integration of automated feature engineering principles into dynamic multi-objective optimization for drug discovery, extending beyond traditional symbolic regression approaches; (2) a semi-automated framework (AMODO-EO) that systematically discovers and validates chemically meaningful objectives during optimization while maintaining performance; (3) experimental validation on ChEMBL datasets demonstrating consistent discovery of interpretable objectives (e.g., the HBA/RTB ratio) with rigorous statistical validation; and (4) comprehensive sensitivity analysis and ablation studies quantifying algorithmic robustness. We show that discovered objectives meet multi-level validity checks (statistical independence, discriminative power, chemical interpretability) and provide actionable design insights. By extending multi-objective formulations beyond fixed criteria through domain-aware objective discovery, AMODO-EO opens a new research direction for adaptive optimization in drug design.

\section{AMODO-EO: Objective Discovery Framework}

\subsection{Problem Formulation}

Let $\mathcal{X} \subset \mathbb{R}^n$ be the feasible chemical space, where each $x \in \mathcal{X}$ encodes a molecule by $n$ descriptors (e.g., molecular weight, LogP, etc.). The traditional multi-objective drug optimization problem is:

\begin{equation}
\min_{x \in \mathcal{X}} \mathbf{F}(x) = [f_1(x), f_2(x), \dots, f_k(x)]^T
\end{equation}

subject to constraints $g(x)\le 0$, $h(x)=0$. Here each $f_i: \mathcal{X}\to \mathbb{R}$ is a user-defined objective (e.g., binding affinity, drug-likeness composite score, synthetic accessibility). This fixed-objective formulation assumes all relevant criteria are known in advance.

AMODO-EO generalizes this to allow emergent objectives that can appear during optimization. At iteration $t$, let

\begin{equation}
\mathbf{F}^t(x) = [f_1(x), \dots, f_k(x), e^{(t)}_{1}(x), \dots, e^{(t)}_{m_t}(x)]^T,
\end{equation}

where $e^{(t)}_j(x)$ are newly discovered objective functions (based on descriptor combinations) up to time $t$. Initially $m_0=0$. Our goal is to identify valid new objectives from data and add them to the optimization.
\begin{definition}[Emergent Objective]
A function $e:\mathcal{X}\to \mathbb{R}$ discovered during optimization is accepted as a new objective if it meets the following criteria:
\begin{enumerate}
    \item \textbf{Mathematical Validity:} $e(x)$ is well-defined (e.g., no division-by-zero) over $\mathcal{X}$.
    \item \textbf{Statistical Independence:} Its values over the current population have low Pearson correlation $|\rho(e, f_i)| < \rho_{\text{thresh}}$ (we use $\rho_{\text{thresh}}=0.7$) with all existing objectives $f_i$.
    \item \textbf{Discriminative Power:} $\mathrm{Var}(e(\mathcal{P}_t)) > \text{var}_{\text{thresh}}$ (we use $\text{var}_{\text{thresh}} = 0.01\times\text{mean}(\mathrm{Var}(f_i(\mathcal{P}_t)))$), ensuring $e$ is not nearly constant.
    \item \textbf{Chemical Interpretability:} The functional form of $e$ corresponds to a meaningful descriptor relationship, validated through a combination of automated pre-filtering (rejecting known unstable combinations, e.g., LogP in denominators) and domain expert assessment for chemical relevance. This semi-automated approach balances computational efficiency with chemical validity, ensuring discovered objectives are scientifically meaningful rather than purely statistical artifacts.
\end{enumerate}
\end{definition}

Criteria (2)–(3) ensure statistical independence via Pearson correlation significance (null $H_0: \rho=0$) and require $e$ to capture non-trivial variance. The interpretability check represents a deliberate design choice to maintain chemical grounding: while this introduces a human-in-the-loop component that limits full automation, it prevents the acceptance of statistically valid but scientifically meaningless relationships. The automated pre-filtering component rejects combinations known to be chemically problematic (e.g., LogP in denominators for hydrophilic compounds, rotatable bonds in denominators due to division-by-zero risks), while expert validation ensures alignment with established chemical principles. Future implementations could enhance automation through integration with chemical ontology systems, literature mining, or knowledge-based scoring functions while preserving chemical relevance.

\subsection{Objective Discovery Operator}

We define an objective discovery operator $\Phi(\mathcal{S},H)$ that maps the current solution set $\mathcal{S}$ and historical data $H$ to a set of candidate objectives. In practice, at each generation we enumerate functions of pairs of descriptors using predefined forms:

\begin{align}
f_{\text{ratio}}(d_i,d_j) &= \frac{d_i}{d_j + \delta}, \\
f_{\text{product}}(d_i,d_j) &= d_i \cdot d_j, \\
f_{\text{diff}}(d_i,d_j) &= |d_i - d_j|,
\end{align}

where $\delta=1$ is a small constant avoiding division by zero. These cover many intuitive relationships (e.g., HBD/(HBA+1), MW–TPSA difference, etc.). To discover objectives involving more than two descriptors, we also allow iterative composition: once a new candidate is validated and added, it becomes available as a descriptor in future candidate generation. We limit the composition depth to 2 levels (i.e., functions of functions) and consider at most 3 descriptors per composite objective to maintain interpretability and tractability. 

For example, the discovered \emph{Selectivity Score} objective $c(x)= \frac{\mathrm{HBD}(x)}{\mathrm{HBA}(x)+1}\times \mathrm{MW}(x)$ arises by first forming the ratio $\mathrm{HBD}/(\mathrm{HBA}+1)$ and then multiplying by MW. Each generated candidate $c$ is then subjected to the validation checks above: we compute its Pearson correlation with all current objectives and its variance over the population $\mathcal{P}_t$. We retain $c$ only if $|\rho(c,f_i)|<\rho_{\text{thresh}}$ for all $i$ and $\mathrm{Var}(c) > \text{var}_{\text{thresh}}$. Finally, we perform a chemical interpretability check: for instance, a ratio of two H-bond descriptors is chemically sensible, while an arbitrary combination of unrelated descriptors would be discarded. Validated candidates are then added: $\mathbf{F}^{t+1} = \mathbf{F}^t \cup \{c\}$.

\textbf{Control of Objective Growth:} The thresholds on correlation and variance (along with the depth limit) control the growth of objectives. New objectives are generated at every generation from all eligible descriptor pairs, but only those passing the strict independence and variance criteria are added. In practice, this results in only a few emergent objectives per run (typically on the order of 2–3), preventing unbounded growth. If no candidate passes the filters in a given generation, the objective set remains unchanged. This dynamic generation process is depicted in Figure~1, which schematically illustrates the AMODO-EO pipeline (objective discovery, filtering, and integration in each generation).

\subsection{Adaptive Weight Evolution}

We maintain a weight vector $\mathbf{w}$ of length $k + m_t$ that represents the relative importance of each objective in guiding the search. These weights are adapted to steer the population toward preferred trade-offs. Concretely, we use a loss function combining a weighted hypervolume objective, an $L_2$ norm regularizer, and an entropy term for diversity:

\begin{equation}
L(\mathbf{w}, \mathcal{P}_t) = -\mathrm{HVI}_w(\mathcal{P}_t) + \lambda\|\mathbf{w}\|_2^2 + \mu \sum_i H(w_i),
\end{equation}

where $\mathrm{HVI}_w(\mathcal{P}_t)$ is the hypervolume indicator of the current population after scaling each objective by its weight $w_i$ (i.e.\ a weighted hypervolume), $\lambda$ is a regularization factor, and $H(w_i) = -w_i\log(w_i + \epsilon)$ is the Shannon entropy (with small $\epsilon$ for stability). Maximizing the hypervolume encourages selection of weights that improve Pareto-front coverage, while the entropy term discourages concentrating all weight on a single objective, promoting a more uniform preference spread.

To update $\mathbf{w}$, we compute the gradient $\nabla_{\mathbf{w}} L$. In effect, this requires the gradient of the (weighted) hypervolume term with respect to $w$. Prior work has shown that the gradient of hypervolume can be computed (e.g., using an uncrowded indicator) and is equivalent to a self-adjusting weighted sum of individual objective gradients. In practice, we approximate $\nabla_{\mathbf{w}}(-\mathrm{HVI}_w)$ numerically via finite differences: for example, perturbing each $w_i$ by a small $\delta$ and observing the change in the scalarized optimization outcome (since exact differentiation through the hypervolume is complex, as the hypervolume is non-convex and piecewise-smooth). Thus $\nabla_{\mathbf{w}}L$ is computed approximately, and we perform a gradient step $\mathbf{w}_{\text{grad}} = \mathbf{w} - \alpha \nabla_{\mathbf{w}}L$ with learning rate $\alpha$.

The entropy regularization $\mu \sum_i H(w_i)$ encourages a diverse weight distribution: maximizing entropy pushes toward a uniform vector, avoiding degenerate solutions that focus on a single objective. We find this helpful in high-dimensional objective spaces to keep the search well-rounded.

\paragraph{Role of Weights in Search.} It is important to clarify that in AMODO-EO the weight vector $\mathbf{w}$ does not directly rank or filter the solutions during selection. We continue to use standard Pareto dominance (nondominated sorting) on the objective set $\mathbf{F}^t$ for selection. Instead, the weights influence the search indirectly: they adjust the focus of the optimization by weighting the hypervolume measure and thus indirectly affecting which trade-offs are favored. In other words, $\mathbf{w}$ guides the evolution of preferences, but does not alter the dominance relation in NSGA-II. The result is that the Pareto-based selection still uses all objectives equally, while the weight update drives the search toward regions that improve the (weighted) hypervolume.

\subsection{Conflict Resolution Matrix}

Because objectives can be redundant or conflicting, we introduce a conflict-resolution step. We compute a conflict matrix $C \in \mathbb{R}^{(k+m_t)\times(k+m_t)}$ whose entry $C_{ij}$ quantifies the conflict intensity between objectives $i$ and $j$. We set

\begin{equation}
C_{ij} = -\frac{\mathrm{cov}(f_i(\mathcal{P}_t),\,f_j(\mathcal{P}_t))}{\sqrt{\mathrm{var}(f_i(\mathcal{P}_t))\cdot \mathrm{var}(f_j(\mathcal{P}_t))}},
\end{equation}

i.e.\ the negative of the Pearson correlation between objectives over the current population. Thus, $C_{ij}$ is negative when $f_i$ and $f_j$ are positively correlated (redundant) and positive when they are negatively correlated (conflicting). Under this definition, $C_{ij}=0$ when objectives are uncorrelated, and higher positive values indicate stronger conflict. We interpret larger $C_{ij}$ as more conflict (objectives moving in opposite directions).

We then solve a small quadratic program to adjust the weights based on $C$. Specifically, we solve

\begin{equation}
\mathbf{w}^* = \arg\min_{\mathbf{w}} \ \mathbf{w}^T C \,\mathbf{w} \quad \text{subject to } \|\mathbf{w}\|_1 = 1,\ \mathbf{w}\ge 0,
\end{equation}

so that pairs of positively correlated objectives (which produce negative $C_{ij}$) receive lower combined weight. This QP (a constrained convex minimization) encourages the total conflict to be small. In effect, minimizing $\mathbf{w}^T C \mathbf{w}$ drives weights down for objective pairs that are positively correlated (redundant), since those produce negative contributions; it allows more weight on objective pairs that are negatively correlated (high $C_{ij}$), since those terms add positive value. We solve (9) at each iteration to obtain a conflict-adjusted weight vector $\mathbf{w}_{\text{conflict}}$.

Finally, we combine the two weight updates:
\begin{equation}
\mathbf{w}^{t+1} = \beta\,\mathbf{w}_{\text{grad}} + (1-\beta)\,\mathbf{w}_{\text{conflict}},
\end{equation}
with $\beta=0.7$. The parameter $\beta$ balances hypervolume-driven learning (larger $\beta$) versus conflict reduction (smaller $\beta$). We found $\beta\approx0.7$ to work well across datasets, providing a robust balance between the two objectives.

\section{The AMODO-EO Algorithm}

Algorithm~\ref{alg:amodo} summarizes the complete AMODO-EO procedure. We assume an initial population $P_0$ of $N$ molecules (vectors $x\in \mathcal{X}$), randomly sampled to ensure diversity across runs. The initial objective set is $F_0 = \{f_1, \dots, f_k\}$. The main loop runs for $T$ generations. At each generation $t$, we:
\begin{enumerate}
    \item Evaluate $P_t$ on all objectives in $F_t$.
    \item (Optional) Update any surrogate model $\Psi(P_t)$ if used.
    \item \textbf{Objective Discovery:} Generate candidate functions using ratio, product, and difference forms on all eligible descriptor pairs (including new ones up to the depth limit). For each candidate $c$:
    \begin{enumerate}
        \item Compute Pearson correlation $\rho(c,f_i)$ with all existing objectives $f_i\in F_t$.
        \item If $|\rho(c,f_i)| < \rho_{\text{thresh}}$ for all $i$ and $\mathrm{Var}(c) > \text{var}_{\text{thresh}}$, then \textbf{keep} $c$; else discard it.
        \item Perform chemical interpretability check on each kept $c$. If interpretable, add to the set of new objectives.
    \end{enumerate}
    \item Add all validated new objectives to $F_{t}$: $F_{t+1} = F_t \cup \{\text{new }c\}$.
    \item \textbf{Weight Evolution:} Compute $\nabla_{\mathbf{w}}L(\mathbf{w},P_t)$ for loss $L = -\mathrm{HVI}_w(P_t) + \lambda\|\mathbf{w}\|_2^2 + \mu\sum_i H(w_i)$. Update $\mathbf{w}_{\text{grad}} = \mathbf{w} - \alpha\nabla_{\mathbf{w}}L$.
    \item \textbf{Conflict Resolution:} Update conflict matrix $C_{t+1}$ using (8). Solve the QP (9) to obtain $\mathbf{w}_{\text{conflict}}$.
    \item \textbf{Combine Updates:} $\mathbf{w} \leftarrow \beta\,\mathbf{w}_{\text{grad}} + (1-\beta)\,\mathbf{w}_{\text{conflict}}$.
    \item \textbf{Selection/Reproduction:} Perform a nondominated sort of $P_t$ using objectives in $F_{t+1}$, then generate offspring via crossover and mutation. Form $P_{t+1}$ by selecting the best $N$ individuals (e.g., by crowding distance). 
    \item Update history: $H_{t+1} = H_t \cup \{(P_t, F_t, \mathbf{w})\}$.
\end{enumerate}
The algorithm then returns the final population $P_T$ and discovered objectives $E^*$. This high-level procedure ensures precise handling of discovered objectives. The weight vector $\mathbf{w}$ guides the focus of the search but does not alter the Pareto sorting (which remains on all objectives). In practice, we find that every new objective added causes the Pareto set to adapt and explore along that new dimension, while still covering the original objectives.

\begin{algorithm}[h]
\caption{AMODO-EO (Emergent Objective Discovery MOO)}
\label{alg:amodo}
\begin{algorithmic}[1]
\REQUIRE Initial population $P_0$ (random, size $N$), initial objectives $F_0=\{f_1,\dots,f_k\}$, parameters $\{\alpha,\beta,\lambda,\mu,\rho_{\text{thresh}},\text{var}_{\text{thresh}}\}$.
\ENSURE Final population $P^*$, discovered objectives $E^*$.
\STATE Initialize weights $\mathbf{w}_0 = \frac{1}{k}\mathbf{1}$ (uniform), history $H_0 = \emptyset$.
\FOR{$t=0$ to $T-1$}
  \STATE Evaluate $P_t$ on all objectives in $F_t$.
  \STATE {\bf Objective Discovery:} 
    \STATE \quad Generate candidates from all descriptor pairs.
    \FOR{each candidate $c$}
      \STATE \quad\quad Compute $\rho(c,f_i)$ for all $f_i\in F_t$.
      \STATE \quad\quad \textbf{if} $|\rho(c,f_i)| < \rho_{\text{thresh}}\ \forall i$ {\bf and} $\mathrm{Var}(c) > \text{var}_{\text{thresh}}$, \textbf{then} keep $c$.
    \ENDFOR
    \STATE \quad Validate interpretability of kept candidates.
    \STATE \quad Add validated candidates to $F_{t}$: $F_{t+1} \gets F_t \cup \{\text{new }c\}$.
  \STATE {\bf Weight Evolution:} Compute $\nabla_{\mathbf{w}}L$ for $L$ in (7); set $\mathbf{w}_{\text{grad}} \gets \mathbf{w} - \alpha \nabla_{\mathbf{w}}L$.
  \STATE {\bf Conflict Resolution:} Update conflict matrix $C_{t+1}$ using (8); solve QP (9) to get $\mathbf{w}_{\text{conflict}}$.
  \STATE {\bf Combine:} $\mathbf{w} \gets \beta \,\mathbf{w}_{\text{grad}} + (1-\beta)\,\mathbf{w}_{\text{conflict}}$.
  \STATE {\bf Selection/Reproduction:} Nondominated sort $P_t$ using $F_{t+1}$; generate offspring via crossover/mutation; select best $N$ to form $P_{t+1}$.
  \STATE Update history: $H_{t+1} \gets H_t \cup \{(P_t, F_t, \mathbf{w})\}$.
\ENDFOR
\STATE \textbf{return} $P_T$, discovered objectives $E^* = F_T \setminus \{f_1,\dots,f_k\}$.
\end{algorithmic}
\end{algorithm}

\section{Experimental Methodology}

\subsection{Datasets and Configuration}
We evaluate AMODO-EO on three datasets of small molecules from the ChEMBL database \citep{gaulton2017chembl,mendez2019chembl}, chosen to represent typical drug-discovery scenarios:
\begin{itemize}
    \item \textbf{Benchmark Dataset:} 400 compounds with 5 molecular descriptors (a simple synthetic test).
    \item \textbf{ChEMBL Small:} 1,000 compounds targeting the dopamine D2 receptor (ChEMBL ID 279).
    \item \textbf{ChEMBL Large:} 5,000 compounds from the same target class, drawn randomly from ChEMBL entries.
\end{itemize}
Each compound is represented by descriptors: molecular weight (MW), logP, hydrogen bond acceptors (HBA), hydrogen bond donors (HBD), topological polar surface area (TPSA), number of rotatable bonds (RTB), and number of aromatic rings. These are common in drug design (e.g., Lipinski’s rule of five involves MW, HBA/HBD, and logP; TPSA and RTB relate to bioavailability).

The initial objectives are:
\begin{enumerate}
    \item \textbf{Binding Affinity:} $f_{\text{bind}}(x) = -\text{pIC}_{50}(x)$, using experimental activity data from ChEMBL (so higher potency means more negative $f_{\text{bind}}$). Here $\mathrm{pIC}_{50} = -\log_{10}(\text{IC}_{50})$ is the potency measure.
    \item \textbf{Drug-likeness Composite Score:} $f_{\text{DL}}(x) = -\left[0.4\,\mathrm{SA}(x) + 0.3\,\mathrm{QED}(x) + 0.3\,\mathrm{Lip}(x)\right]$, where SA is the synthetic accessibility score, QED is the quantitative estimate of drug-likeness (higher is better), and Lip is the Lipinski compliance count (0–5). We take a negative so that higher drug-likeness yields lower objective.
    \item \textbf{Synthetic Accessibility:} $f_{\text{SA}}(x) = \mathrm{SA}(x)$ (we seek to minimize synthetic difficulty).
\end{enumerate}
All objectives are set as minimization problems (we invert signs as needed). For clarity, pIC$_{50}$ is defined as $-\log_{10}(\text{IC}_{50})$, so that larger pIC$_{50}$ indicates higher affinity.

Algorithm parameters are set by preliminary tuning: population size $N=100$, generations $T=50$, and standard NSGA-II crossover/mutation operators \citep{deb2002fast}. We set $\rho_{\text{thresh}}=0.7$ for correlation (Definition 1), variance threshold $0.01\times$ average objective variance, learning rate $\alpha=0.05$, entropy weight $\mu=0.1$, regularization $\lambda=0.01$, and weight-mixing parameter $\beta=0.7$. 

\paragraph{Parameter Selection Rationale:} The correlation threshold of 0.7 was chosen to balance objective independence while allowing for moderately correlated but still informative objectives \citep{warren2006comparative}. This threshold aligns with standard statistical practice where correlations above 0.7 indicate potential redundancy \citep{cross2009comparison}. The population size of 100 and 50 generations provide sufficient sampling and convergence time based on preliminary experiments \citep{zitzler2001spea2}, while remaining computationally feasible for the descriptor space size. The learning rate and regularization parameters were selected to ensure stable weight evolution without over-fitting to short-term hypervolume fluctuations \citep{zhang2007moead}.

These parameters yielded robust performance across all datasets. Each experiment is repeated 5 times with different random seeds (random initial populations) and we report averages and standard deviations to ensure statistical reliability of the results.

\subsection{Evaluation Metrics}
We evaluate performance in terms of (a) objective discovery metrics (number and quality of emergent objectives) and (b) optimization quality (hypervolume of final fronts). For discovery, we define \emph{Discovery Rate} as the fraction of independent runs in which a new objective was found, and \emph{Avg.\ Objectives} as the average number of validated objectives discovered per run. We also report the specific emergent objectives found (e.g.\ HBA/RTB ratio) and their statistics (correlation range, variance, significance). 

\paragraph{Metric Selection Rationale:} The discovery rate provides a measure of the framework's reliability and consistency in finding new objectives across independent runs, which is crucial for establishing the robustness of the discovery mechanism. The correlation and variance statistics for discovered objectives ensure they meet our strict independence and discriminative power criteria, preventing the acceptance of redundant or trivial objectives. 

For optimization quality, we compare the final hypervolume of the Pareto fronts on the original objectives against a baseline NSGA-II run with no emergent objectives. While this comparison has inherent limitations once new objectives are added to the problem space, it provides insight into the computational trade-offs involved in the discovery process and ensures that the framework maintains reasonable performance on the user-defined criteria.

\section{Results and Analysis}

\subsection{Objective Discovery Results}
AMODO-EO consistently discovered several meaningful emergent objectives across datasets. Most notably, the ratio HBA/RTB (hydrogen bond acceptors to rotatable bonds) was found in all 5 independent runs on the ChEMBL datasets, indicating a very robust pattern. Other frequent discoveries included the MW/TPSA ratio, the product of LogP and aromatic ring count, and composite functions such as the \emph{Selectivity Score} and \emph{Novelty Metric} defined below. These emergent objectives passed the statistical filters ($|\rho|<0.7$) and had significant variance (Table~\ref{tab:validation}).

\begin{table}[h]
\centering
\caption{Emergent objectives discovered by AMODO-EO across datasets.}
\label{tab:objectives}
\begin{tabular}{l|l|c}
\toprule
\textbf{Dataset} & \textbf{Discovered Objective} & \textbf{Discovery Frequency} \\
\midrule
Benchmark    & MW-TPSA ratio         & 4/5 runs \\
ChEMBL Small & HBA/RTB ratio         & 5/5 runs \\
ChEMBL Large & LogP$\times$(aromatic rings) & 3/5 runs \\
\bottomrule
\end{tabular}
\end{table}

\subsubsection{Case Study – HBA/RTB Ratio}
In the ChEMBL Small dataset, the HBA/RTB ratio consistently emerged as a chemically meaningful objective. This ratio is defined as the number of hydrogen bond acceptors (HBA) divided by (RTB$+1$) for regularization. It reflects the balance between molecular polarity (HBA) and conformational flexibility (RTB), a trade-off known to influence potency, permeability, and overall drug-likeness.

\textbf{Statistical Validation:} The HBA/RTB ratio exhibited Pearson correlation coefficients ranging from $-0.12$ to $0.31$ with the existing objectives (binding affinity, drug-likeness score, synthetic accessibility), all well below the 0.7 independence threshold. Its variance across the population was 0.087, exceeding the discriminative power threshold (Table~\ref{tab:validation}), confirming that it differentiates candidates.

\textbf{Chemical Interpretation:} Medicinal chemistry literature extensively documents the relationship between rotatable bonds and molecular flexibility, membrane permeability \citep{veber2002molecular,egan2000prediction}, while hydrogen bond acceptors influence solubility and target binding \citep{lipinski2001experimental,muegge2001simple}. Higher HBA/RTB ratios suggest molecules with strong hydrogen-bonding potential relative to their flexibility, correlating with improved binding affinity and favorable drug-likeness \citep{hughes2011physiochemical,gleeson2008generation}. This emergent objective captures a design trade-off that was not explicitly specified a priori but aligns with established structure-activity relationships in medicinal chemistry \citep{ghose1999knowledge,oprea2000property}.

\textbf{Discovery Performance:} The HBA/RTB objective was detected in 5/5 independent runs on the ChEMBL dataset, demonstrating robust detectability by the algorithm’s discovery mechanism. When included in optimization, the Pareto front expanded along this new dimension: solutions emerged with higher HBA relative to RTB, reflecting the learned trade-off. This extended the diversity of solutions while preserving coverage of the original objectives (see Section~5.3).

\subsubsection{Other Discovered Objectives}
In addition to HBA/RTB, the algorithm uncovered several other chemically interpretable objectives:
\begin{itemize}
    \item \textbf{Selectivity Score:} Discovered in 3/5 runs, defined as 
    \begin{equation}
        f_{\text{sel}}(x) = \frac{\mathrm{HBD}(x)}{\mathrm{HBA}(x) + 1} \times \mathrm{MW}(x).
    \end{equation}
    This combines hydrogen-bond donors and acceptors with molecular size, aiming to capture specificity between target and off-target receptors. Its Pearson correlations with existing objectives were in $[-0.15,0.28]$, variance 0.072 (above threshold), and it passed independence tests ($p<0.05$).
    \item \textbf{Synthetic Complexity:} Found in 4/5 runs, defined as
    \begin{equation}
        f_{\text{syn}}(x) = \mathrm{RTB}(x)\cdot \mathrm{AromaticRings}(x) + \frac{\mathrm{TPSA}(x)}{100}.
    \end{equation}
    This blends flexibility, aromaticity, and polarity; higher values suggest more complex synthesis. It had correlations $[0.22,0.31]$ (positive, low redundancy) and variance 0.051.
    \item \textbf{Novelty Metric:} In 2/5 runs, defined as
    \begin{equation}
        f_{\text{nov}}(x) = \left|\frac{\log P(x) - \mathrm{RTB}(x)}{5}\right|.
    \end{equation}
    This measures deviation from an expected lipophilicity–flexibility relationship, indicating how “outlier” a molecule is in chemical space. The rationale is that typical drug-like molecules follow an empirical relationship where LogP scales roughly with RTB/5. Larger deviations suggest novel structures. Its correlation range was $[0.09,0.24]$, variance 0.038 (above threshold), and significance $p<0.10$.
\end{itemize}
All emergent objectives above passed the independence and variance checks (see Table~\ref{tab:validation}), confirming they are not artifacts of the original design criteria but true discoveries from systematic search.

\subsection{Ablation Study}
To quantify the importance of each component, we performed an ablation study (Table~\ref{tab:ablation}). We measured the optimization performance (final hypervolume) under configurations: (a) full AMODO-EO, (b) without objective discovery (baseline NSGA-II), (c) without statistical filtering (accepting all generated candidates), and (d) without chemical interpretability checks (accepting statistically valid but potentially meaningless objectives). The results demonstrate that the statistical filtering components are crucial for ensuring validity, while the interpretability filter prevents acceptance of spurious relationships that could mislead the optimization process.

\begin{table}[h]
\centering
\caption{Ablation study: component contributions to optimization performance.}
\label{tab:ablation}
\begin{tabular}{lcc}
\toprule
Component Configuration & Final Hypervolume & Performance Change \\
\midrule
Full AMODO-EO                 & $0.823 \pm 0.029$ & Baseline \\
Without Objective Discovery   & $0.847 \pm 0.023$ & $+2.9\%$ \\
Without Statistical Filtering & $0.798 \pm 0.035$ & $-3.0\%$ \\
Without Interpretability      & $0.801 \pm 0.032$ & $-2.7\%$ \\
\bottomrule
\end{tabular}
\end{table}

\subsection{Performance Trade-off Analysis}
We evaluated the optimization performance of AMODO-EO, with a focus on understanding the exploration-exploitation trade-offs inherent in objective discovery. Rather than using the methodologically problematic approach of comparing cross-dimensional hypervolumes, we analyze the performance implications more appropriately.

\textbf{Performance on Original Objectives:} Table~4 shows the final hypervolume of AMODO-EO compared to baseline NSGA-II, both evaluated on the original three objectives. AMODO-EO shows a modest reduction in hypervolume (2.4\% on Small, 1.5\% on Large), which should be interpreted as the computational cost of discovery rather than a performance deficit. The algorithm allocates a portion of its search budget to exploring potential objectives rather than exclusively exploiting known ones.

\textbf{Solution Diversity Analysis:} AMODO-EO generates Pareto fronts that span higher-dimensional spaces when discovered objectives are included. While direct hypervolume comparison across different dimensionalities is not meaningful, qualitative analysis shows that the original objective trade-offs are largely preserved while new design dimensions are revealed. The discovered objectives provide orthogonal design axes that would be invisible to traditional fixed-objective optimization.

\textbf{Design Space Coverage:} The inclusion of discovered objectives like HBA/RTB ratio reveals solution clusters with distinct chemical profiles that would not emerge in traditional optimization. This expansion of the design space provides chemists with a more complete view of molecular trade-offs, representing valuable scientific insights that justify the modest performance cost on original objectives.

\begin{table}[h]
\centering
\caption{Optimization performance (hypervolume on original objectives).}
\label{tab:performance}
\begin{tabular}{l|c|c}
\toprule
\textbf{Algorithm} & \textbf{ChEMBL Small} & \textbf{ChEMBL Large} \\
\midrule
NSGA-II (fixed objectives)    & $0.847\pm0.023$ & $0.834\pm0.031$ \\
AMODO-EO (with discovery)     & $0.823\pm0.029$ & $0.819\pm0.027$ \\
\midrule
\textbf{Performance Cost}     & $2.9\%$ & $1.8\%$ \\
\bottomrule
\end{tabular}
\end{table}

The results demonstrate that AMODO-EO successfully balances exploitation of known objectives with exploration of new ones, yielding actionable chemical insights at a modest computational cost.

\subsection{Statistical Validation}
Table~\ref{tab:validation} summarizes statistical validation of all discovered objectives (aggregated across datasets). Discovery Rate here means the fraction of runs in which that specific objective was found (e.g.\ 5/5 for HBA/RTB). We report the range of correlation coefficients with existing objectives, the objective’s variance, and the $p$-value for the null hypothesis of zero correlation. All discovered objectives have $|\rho|<0.7$ by design and pass significance tests (typically $p<0.05$, except Novelty uses $p<0.10$ due to fewer occurrences). The HBA/RTB ratio stands out with 5/5 runs and $p<0.01$. These statistics ensure the objectives are neither redundant nor trivial.

\begin{table}[h]
\centering
\caption{Statistical validation of all discovered objectives (ChEMBL datasets).}
\label{tab:validation}
\begin{tabular}{l|c|c|c|c}
\toprule
\textbf{Objective} & \textbf{Discovery Rate} & \textbf{Corr.\ Range} & \textbf{Variance} & \textbf{$p$-value} \\
\midrule
HBA/RTB ratio           & 5/5 runs & [0.12, 0.31] & 0.087 & $<0.01$ \\
MW/TPSA ratio          & 4/5 runs & [0.18, 0.25] & 0.043 & $<0.05$ \\
LogP–Aromatic product  & 3/5 runs & [0.21, 0.33] & 0.058 & $<0.05$ \\
Selectivity Score      & 3/5 runs & [0.15, 0.28] & 0.072 & $<0.05$ \\
Synthetic Complexity   & 4/5 runs & [0.22, 0.31] & 0.051 & $<0.01$ \\
Novelty Metric         & 2/5 runs & [0.09, 0.24] & 0.038 & $<0.10$ \\
\bottomrule
\end{tabular}
\end{table}

In summary, AMODO-EO reliably uncovers multiple emergent objectives that meet our validity criteria, demonstrating systematic discovery capability. In particular, the HBA/RTB ratio, consistently found in all runs, highlights a genuine structural pattern (balancing polarity and flexibility) in drug-like molecules. The combination of filter thresholds and interpretability ensures these objectives provide chemically meaningful axes for design beyond the original objectives.

\subsection{Sensitivity Analysis}
To address concerns about parameter dependence and demonstrate algorithmic robustness, we conducted comprehensive sensitivity analysis on key algorithmic parameters. The results confirm that AMODO-EO maintains consistent discovery performance across reasonable parameter ranges.

\textbf{Correlation Threshold ($\rho_{\text{thresh}}$):} Varying from 0.5 to 0.9, we observed discovery rate changes of $\pm 12\%$. The HBA/RTB objective remained discoverable across this entire range, with optimal performance at $\rho_{\text{thresh}} = 0.7$. Lower thresholds (0.5-0.6) permitted more redundant objectives, while higher thresholds (0.8-0.9) were overly restrictive, filtering out potentially valuable relationships.

\textbf{Weight-Mixing Parameter ($\beta$):} Testing $\beta \in [0.5, 0.9]$ showed discovery rate variations of only $\pm 3\%$. The algorithm demonstrates remarkable stability to this parameter, with $\beta = 0.7$ providing optimal balance between hypervolume-driven and conflict-driven weight updates.

\textbf{Learning Rate ($\alpha$):} Varying from 0.01 to 0.1 showed $\pm 8\%$ discovery rate changes. Values below 0.03 resulted in slow convergence of weight updates, while values above 0.08 caused instability in the gradient approximation. The chosen value of 0.05 provides stable gradient-based learning without oscillations.

\textbf{Variance Threshold Multiplier:} Testing multipliers from 0.005 to 0.02 showed $\pm 6\%$ discovery rate changes. The chosen value of 0.01 effectively filters trivial, near-constant objectives while retaining those with meaningful discriminative power.

These results confirm that the framework's core discovery capability is not critically dependent on precise parameterization. Most importantly, the signature HBA/RTB objective was consistently discoverable across all tested parameter ranges, demonstrating that the fundamental discovery mechanism identifies robust chemical relationships rather than parameter-dependent artifacts.

\section{Discussion}
AMODO-EO advances the state-of-the-art by automating the identification of new objectives in molecular optimization. Prior dynamic MOO methods \citep{jin2005evolutionary,deb2002fast} adapt to changing objectives only when these are externally specified. In contrast, our framework discovers objectives endogenously from the data structure, representing a significant advancement over traditional approaches such as fixed-objective NSGA-II \citep{deb2002fast}, SPEA2 \citep{zitzler2001spea2}, and MOEA/D \citep{zhang2007moead}.

\textbf{Comparison with Existing Approaches:} Traditional multi-objective optimization methods in drug discovery \citep{brown2019guacamol,polykovskiy2020molecular} rely on predetermined objective functions, typically including binding affinity prediction \citep{wang2003further,cheng2009comparative}, drug-likeness scores \citep{bickerton2012quantifying,gleeson2008generation}, and synthetic accessibility \citep{ertl2009estimation}. While these approaches have been successful in specific applications \citep{segler2018planning,jin2018junction}, they cannot adapt to discover new optimization criteria during the search process.

Recent advances in automated molecular design \citep{gomez2018automatic,schwaller2019molecular,winter2019learning} have focused on generative models and representation learning, but have not addressed the fundamental limitation of fixed objective sets. Our approach complements these advances by providing a systematic framework for objective discovery that can be integrated with any underlying molecular representation or generation method.

The main contributions relative to prior work are: (i) a systematic objective discovery mechanism, extending dynamic MOO to search over possible descriptor functions; (ii) a statistical validation framework for filtering emergent objectives based on independence and variance; (iii) integration of discovered objectives with adaptive weighting and conflict resolution; and (iv) empirical demonstration that discovered objectives (such as HBA/RTB) provide new, interpretable design criteria. This fills a gap in multi-objective optimization for drug design by not assuming that all relevant objectives are known a priori \citep{schneider2018automating,clark2014drug}.

\subsection{The Cost of Discovery: Interpreting Performance Trade-offs}
The modest reduction in hypervolume observed when AMODO-EO is compared to traditional NSGA-II on the original objective space should be understood as the necessary cost of scientific discovery rather than a performance limitation. Traditional optimization algorithms dedicate their entire computational budget to exploiting known objectives, while AMODO-EO strategically allocates a portion of its resources to exploring the space of potential new objectives. This exploration-exploitation trade-off is fundamental to any discovery process and represents a conscious design choice to prioritize knowledge acquisition alongside optimization performance. The resulting Pareto front spans a higher-dimensional space that reveals previously hidden trade-offs, providing chemists with a more complete understanding of the molecular design landscape. In this context, the slight reduction in performance on the original objectives represents a valuable investment that yields new, actionable scientific insights.

\subsection{Practical Implications of Objective Discovery}
The objectives found by AMODO-EO offer actionable chemical insights. For instance, the HBA/RTB ratio reveals that successful D2 ligands tend to trade off hydrogen bonding capacity against molecular flexibility. Chemically, this suggests that adding too many rotatable bonds (increasing flexibility) must be balanced by polar features (acceptors) to maintain potency. Such a relationship was not encoded in the original objectives, yet the algorithm uncovered it consistently, indicating it is a real pattern in the data rather than noise. Similarly, the Selectivity Score and Synthetic Complexity metrics suggest new angles: the former encodes specificity between receptor vs.\ off-target binding, and the latter integrates multiple factors affecting synthetic difficulty. In practice, drug designers could use these emergent criteria to bias compound enumeration: for example, enforcing a minimum HBA/RTB ratio or a maximum synthetic complexity when generating libraries.

\paragraph{Prospective Application Example:} Consider a practical scenario where AMODO-EO discovers the HBA/RTB ratio as a key emergent objective for a dopamine D2 receptor project. Upon this discovery, a design team could immediately implement a new filtering rule within their virtual screening or library enumeration workflow. Specifically, they could prioritize any generated compound where the value of HBA(x)/(RTB(x)+1) exceeds a threshold derived from the Pareto front analysis—for instance, compounds with ratios above 0.5 if the discovered front shows improved activity at this level. This action would directly enrich the subsequent pool of candidate molecules with structures possessing a more favorable, data-driven balance of polarity and flexibility. The team could then synthesize and test a focused library based on this criterion, representing a direct translation of the computational discovery into experimental design decisions.

In essence, AMODO-EO provides a form of automated hypothesis generation. It extends beyond black-box optimization by surfacing descriptor relationships that chemists can interpret. Unlike fixed-objective methods, it adapts the optimization goals in response to the discovered data structure. This adaptive capability is analogous to a human-in-the-loop insight process, but done systematically and at scale.

\subsection{Limitations and Future Work}
We acknowledge several important limitations that define the scope and applicability of our approach.

\textbf{Theoretical Convergence:} Formal convergence analysis remains an open challenge for dynamic multi-objective systems \citep{jin2005evolutionary,zhang2007moead}. The continuously evolving objective set means that classical convergence theorems (which assume fixed Pareto fronts) do not directly apply. Rigorous theoretical analysis would require strong assumptions about the discovery process and objective emergence patterns that may not hold in practice.

\textbf{Discovery Mechanism Limitations:} Our current implementation uses predefined functional forms (ratio, product, difference) limited to pairwise descriptor interactions. This constrains the diversity of discoverable relationships compared to more sophisticated approaches from the symbolic regression literature \citep{schmidt2009distilling,udrescu2020ai}. Grammar-based genetic programming \citep{koza1992genetic} or neural symbolic methods \citep{cranmer2020discovering} could explore richer function spaces, potentially discovering more complex but meaningful chemical relationships.

\textbf{Semi-Automated Interpretability:} The chemical interpretability filter represents a deliberate trade-off between automation and domain validity. While we have automated many aspects (pre-filtering unstable combinations, statistical validation), the final interpretability assessment involves domain expert judgment. This introduces potential subjectivity and limits scalability \citep{bajorath2002integration,schneider2018automating}. However, we argue this is essential for ensuring discovered objectives are scientifically meaningful rather than purely statistical artifacts. Future work should focus on developing automated interpretability scoring through cheminformatics knowledge bases \citep{weininger1988smiles}, literature mining \citep{swanson1986fish}, or ontology-based validation systems.

\textbf{Parameter Sensitivity:} Although our sensitivity analysis demonstrates robustness across reasonable parameter ranges, the framework relies on several heuristically set thresholds. Different chemical domains or optimization contexts may require parameter adaptation. The correlation threshold of 0.7, while grounded in statistical practice \citep{warren2006comparative}, may need adjustment for highly correlated descriptor sets or when stricter independence is required.

\textbf{Limited Validation Scope:} Our experimental validation focuses on one target class (dopamine D2 receptor) and ChEMBL datasets \citep{gaulton2017chembl}. Broader validation across diverse therapeutic targets \citep{overington2006many,hopkins2002druggable}, chemical libraries \citep{sterling2015zinc,irwin2020zinc20}, and optimization contexts is needed to establish generalizability. The discovered relationships (particularly HBA/RTB) may be specific to dopamine receptor ligands and might not generalize to other protein families or chemical scaffolds.

\textbf{Scalability Considerations:} The current implementation evaluates all possible descriptor pairs at each generation, resulting in quadratic computational complexity in the number of descriptors. For high-dimensional descriptor spaces (e.g., fingerprint-based features \citep{rogers2010extended}), more efficient candidate generation strategies would be necessary.

Future directions include: (1) extending validation to broader chemical spaces and diverse therapeutic targets \citep{rask2001genome,imming2006drugs}, (2) integrating more sophisticated discovery mechanisms from the symbolic regression literature \citep{cranmer2020discovering}, (3) developing automated interpretability assessment systems using chemical ontologies \citep{hastings2013chebi}, (4) theoretical analysis of convergence properties under objective discovery, and (5) hybrid approaches combining AMODO-EO with high-fidelity molecular simulations \citep{shirts2003equilibrium,bennett1976efficient} or machine learning models \citep{winter2019learning,schwaller2019molecular}.

\section{Conclusion}
We have presented AMODO-EO, a semi-automated framework that integrates automated feature engineering principles with dynamic multi-objective optimization for drug discovery. Building upon established techniques from symbolic regression and automated feature engineering, our approach addresses the novel challenge of discovering chemically meaningful objectives during active multi-objective optimization while maintaining competitive performance.

The framework successfully demonstrates systematic identification of interpretable molecular relationships, particularly the HBA/RTB ratio that consistently emerged across all experimental runs. This objective captures a genuine chemical trade-off between hydrogen bonding capacity and molecular flexibility that was not explicitly specified in the original optimization formulation. The discovery of such relationships represents valuable chemical insights that can directly inform molecular design strategies.

Our experimental validation on ChEMBL datasets confirms that AMODO-EO maintains competitive optimization performance on original objectives while revealing new design dimensions. The modest performance cost (2-3% hypervolume reduction) represents a reasonable investment for acquiring actionable chemical knowledge that would be invisible to traditional fixed-objective optimization.

The comprehensive sensitivity analysis demonstrates that the framework's core discovery capabilities are robust to parameter variations, with the signature HBA/RTB objective being consistently discoverable across all tested parameter ranges. This robustness indicates that the discovered relationships reflect genuine chemical patterns rather than algorithmic artifacts.

While acknowledging limitations in theoretical convergence guarantees, discovery mechanism sophistication, and validation scope, AMODO-EO opens a new research direction that bridges computational optimization with scientific discovery. By transforming multi-objective optimization from a purely exploitative process into one that systematically explores the space of potential optimization criteria, the framework represents a meaningful advancement in adaptive optimization for chemical design.

The approach's semi-automated nature, incorporating domain expertise for chemical interpretability, represents a practical balance between computational efficiency and scientific validity. This positions AMODO-EO as a valuable tool for computational chemists seeking to uncover hidden molecular trade-offs while maintaining chemical relevance in the optimization process.

\section*{Acknowledgments}
The authors would like to express their sincere gratitude to Mrs. Shwetha S. Shetty, Assistant Professor, and Dr. Rithesh Pakkala P., Associate Professor \& Head, Department of Information Science and Engineering, Sahyadri College of Engineering and Management, for their valuable guidance and support. The authors also thank Dr. Sudheer Shetty, Professor \& Vice Principal, and Dr. Shamantha Rai B, Professor \& Dean-Academic, for their encouragement and academic insights. They are grateful to Dr. S. S. Injaganeri, Principal, for providing excellent facilities and a research-conducive environment. The authors also acknowledge Sahyadri College of Engineering and Management, Mangaluru, for fostering innovation and providing the infrastructure required for this work. Special thanks are extended to Mr. Dinesh Naik, Assistant Professor, Department of Information Technology, NITK Surathkal, for his inspiration and motivation in developing this model.

\begin{thebibliography}{99}
\bibitem{schmidt2009distilling} Schmidt, M. \& Lipson, H. Distilling free-form natural laws from experimental data. \emph{Science} \textbf{324}, 81--85 (2009).

\bibitem{udrescu2020ai} Udrescu, S.-M. \& Tegmark, M. AI Feynman: A physics-inspired method for symbolic regression. \emph{Science Advances} \textbf{6}, eaay2631 (2020).

\bibitem{khurana2018feature} Khurana, U., Samulowitz, H. \& Turaga, D. Feature engineering for predictive modeling using reinforcement learning. \emph{Proceedings of the AAAI Conference on Artificial Intelligence} \textbf{32} (2018).

\bibitem{toyao2020machine} Toyao, T. et al. Machine learning for catalyst informatics: recent applications and prospects. \emph{ACS Catalysis} \textbf{10}, 2260--2297 (2020).

\bibitem{yang2019auto} Yang, K. et al. Analyzing learned molecular representations for property prediction. \emph{Journal of Chemical Information and Modeling} \textbf{59}, 3370--3388 (2019).

\bibitem{deb2002fast} Deb, K., Pratap, A., Agarwal, S. \& Meyarivan, T. A fast and elitist multiobjective genetic algorithm: NSGA-II. \emph{IEEE Transactions on Evolutionary Computation} \textbf{6}, 182--197 (2002).

\bibitem{zitzler2001spea2} Zitzler, E., Laumanns, M. \& Thiele, L. SPEA2: Improving the strength Pareto evolutionary algorithm. \emph{TIK Report} \textbf{103} (2001).

\bibitem{zhang2007moead} Zhang, Q. \& Li, H. MOEA/D: A multiobjective evolutionary algorithm based on decomposition. \emph{IEEE Transactions on Evolutionary Computation} \textbf{11}, 712--731 (2007).

\bibitem{lipinski2001experimental} Lipinski, C. A., Lombardo, F., Dominy, B. W. \& Feeney, P. J. Experimental and computational approaches to estimate solubility and permeability in drug discovery and development settings. \emph{Advanced Drug Delivery Reviews} \textbf{46}, 3--26 (2001).

\bibitem{ertl2009estimation} Ertl, P. \& Schuffenhauer, A. Estimation of synthetic accessibility score of drug-like molecules based on molecular complexity and fragment contributions. \emph{Journal of Cheminformatics} \textbf{1}, 8 (2009).

\bibitem{bickerton2012quantifying} Bickerton, G. R., Paolini, G. V., Besnard, J., Muresan, S. \& Hopkins, A. L. Quantifying the chemical beauty of drugs. \emph{Nature Chemistry} \textbf{4}, 90--98 (2012).

\bibitem{gaulton2017chembl} Gaulton, A. et al. The ChEMBL database in 2017. \emph{Nucleic Acids Research} \textbf{45}, D945--D954 (2017).

\bibitem{mendez2019chembl} Mendez, D. et al. ChEMBL: towards direct deposition of bioassay data. \emph{Nucleic Acids Research} \textbf{47}, D930--D940 (2019).

\bibitem{brown2019guacamol} Brown, N., Fiscato, M., Segler, M. H. \& Vaucher, A. C. GuacaMol: benchmarking models for de novo molecular design. \emph{Journal of Chemical Information and Modeling} \textbf{59}, 1096--1108 (2019).

\bibitem{polykovskiy2020molecular} Polykovskiy, D. et al. Molecular sets (MOSES): a benchmarking platform for molecular generation models. \emph{Frontiers in Pharmacology} \textbf{11}, 565644 (2020).

\bibitem{jin2018junction} Jin, W., Barzilay, R. \& Jaakkola, T. Junction tree variational autoencoder for molecular graph generation. \emph{International Conference on Machine Learning} \textbf{80}, 2323--2332 (2018).

\bibitem{gomez2018automatic} Gómez-Bombarelli, R. et al. Automatic chemical design using a data-driven continuous representation of molecules. \emph{ACS Central Science} \textbf{4}, 268--276 (2018).

\bibitem{segler2018planning} Segler, M. H., Preuss, M. \& Waller, M. P. Planning chemical syntheses with deep neural networks and symbolic AI. \emph{Nature} \textbf{555}, 604--610 (2018).

\bibitem{coley2017prediction} Coley, C. W. et al. Prediction of organic reaction outcomes using machine learning. \emph{ACS Central Science} \textbf{3}, 434--443 (2017).

\bibitem{schwaller2019molecular} Schwaller, P. et al. Molecular transformer: a model for uncertainty-calibrated molecular optimization. \emph{ACS Central Science} \textbf{5}, 1572--1583 (2019).

\bibitem{winter2019learning} Winter, R. et al. Learning continuous and data-driven molecular descriptors by translating equivalent chemical representations. \emph{Chemical Science} \textbf{10}, 1692--1701 (2019).

\bibitem{reymond2015chemical} Reymond, J.-L. The chemical space project. \emph{Accounts of Chemical Research} \textbf{48}, 722--730 (2015).

\bibitem{bohacek1996art} Bohacek, R. S., McMartin, C. \& Guida, W. C. The art and practice of structure-based drug design: a molecular modeling perspective. \emph{Medicinal Research Reviews} \textbf{16}, 3--50 (1996).

\bibitem{kirkpatrick1994exploring} Kirkpatrick, P. \& Ellis, C. Chemical space. \emph{Nature} \textbf{432}, 823 (2004).

\bibitem{fink2007virtual} Fink, T. \& Reymond, J.-L. Virtual exploration of the chemical universe up to 11 atoms of C, N, O, F: assembly of 26.4 million structures (110.9 million stereoisomers) and analysis for new ring systems, stereochemistry, physicochemical properties, and drug discovery. \emph{Journal of Chemical Information and Modeling} \textbf{47}, 342--353 (2007).

\bibitem{ruddigkeit2012enumeration} Ruddigkeit, L., Van Deursen, R., Blum, L. C. \& Reymond, J.-L. Enumeration of 166 billion organic small molecules in the chemical universe database GDB-17. \emph{Journal of Chemical Information and Modeling} \textbf{52}, 2864--2875 (2012).

\bibitem{blum2009970} Blum, L. C. \& Reymond, J.-L. 970 million druglike small molecules for virtual screening in the chemical universe database GDB-13. \emph{Journal of the American Chemical Society} \textbf{131}, 8732--8733 (2009).

\bibitem{sterling2015zinc} Sterling, T. \& Irwin, J. J. ZINC 15--over 750 million compounds for virtual screening. \emph{Journal of Chemical Information and Modeling} \textbf{55}, 2324--2337 (2015).

\bibitem{irwin2020zinc20} Irwin, J. J. et al. ZINC20--a free ultralarge-scale chemical database for ligand discovery. \emph{Journal of Chemical Information and Modeling} \textbf{60}, 6065--6073 (2020).

\bibitem{rogers2010extended} Rogers, D. \& Hahn, M. Extended-connectivity fingerprints. \emph{Journal of Chemical Information and Modeling} \textbf{50}, 742--754 (2010).

\bibitem{morgan1965generation} Morgan, H. L. The generation of a unique machine description for chemical structures--a technique developed at chemical abstracts service. \emph{Journal of Chemical Documentation} \textbf{5}, 107--113 (1965).

\bibitem{weininger1988smiles} Weininger, D. SMILES, a chemical language and information system. 1. Introduction to methodology and encoding rules. \emph{Journal of Chemical Information and Computer Sciences} \textbf{28}, 31--36 (1988).

\bibitem{kramer2012evaluation} Kramer, C., Kalliokoski, T., Gedeck, P. \& Vulpetti, A. The experimental uncertainty of heterogeneous public Ki data. \emph{Journal of Medicinal Chemistry} \textbf{55}, 5165--5173 (2012).

\bibitem{kalliokoski2013comparability} Kalliokoski, T., Kramer, C., Vulpetti, A. \& Gedeck, P. Comparability of mixed IC₅₀ data--a statistical analysis. \emph{PLoS One} \textbf{8}, e61007 (2013).

\bibitem{hughes2011physiochemical} Hughes, J. D. et al. Physiochemical drug properties associated with in vivo toxicological outcomes. \emph{Bioorganic \& Medicinal Chemistry Letters} \textbf{18}, 4872--4875 (2008).

\bibitem{gleeson2008generation} Gleeson, M. P. Generation of a set of simple, interpretable ADMET rules of thumb. \emph{Journal of Medicinal Chemistry} \textbf{51}, 817--834 (2008).

\bibitem{veber2002molecular} Veber, D. F. et al. Molecular properties that influence the oral bioavailability of drug candidates. \emph{Journal of Medicinal Chemistry} \textbf{45}, 2615--2623 (2002).

\bibitem{egan2000prediction} Egan, W. J., Merz, K. M. \& Baldwin, J. J. Prediction of drug absorption using multivariate statistics. \emph{Journal of Medicinal Chemistry} \textbf{43}, 3867--3877 (2000).

\bibitem{muegge2001simple} Muegge, I., Heald, S. L. \& Brittelli, D. Simple selection criteria for drug-like chemical matter. \emph{Journal of Medicinal Chemistry} \textbf{44}, 1841--1846 (2001).

\bibitem{ghose1999knowledge} Ghose, A. K., Viswanadhan, V. N. \& Wendoloski, J. J. A knowledge-based approach in designing combinatorial or medicinal chemistry libraries for drug discovery. 1. A qualitative and quantitative characterization of known drug databases. \emph{Journal of Combinatorial Chemistry} \textbf{1}, 55--68 (1999).

\bibitem{oprea2000property} Oprea, T. I. Property distribution of drug-related chemical databases. \emph{Journal of Computer-Aided Molecular Design} \textbf{14}, 251--264 (2000).

\bibitem{walters1998green} Walters, W. P., Stahl, M. T. \& Murcko, M. A. Virtual screening--an overview. \emph{Drug Discovery Today} \textbf{3}, 160--178 (1998).

\bibitem{shoichet2004virtual} Shoichet, B. K. Virtual screening of chemical libraries. \emph{Nature} \textbf{432}, 862--865 (2004).

\bibitem{kitchen2004docking} Kitchen, D. B., Decornez, H., Furr, J. R. \& Bajorath, J. Docking and scoring in virtual screening for drug discovery: methods and applications. \emph{Nature Reviews Drug Discovery} \textbf{3}, 935--949 (2004).

\bibitem{warren2006essential} Warren, G. L. et al. Essential considerations for using protein-ligand structures in drug discovery. \emph{Drug Discovery Today} \textbf{17}, 1270--1281 (2012).

\bibitem{morris2009automated} Morris, G. M. et al. AutoDock4 and AutoDockTools4: automated docking with selective receptor flexibility. \emph{Journal of Computational Chemistry} \textbf{30}, 2785--2791 (2009).

\bibitem{friesner2004glide} Friesner, R. A. et al. Glide: a new approach for rapid, accurate docking and scoring. 1. Method and assessment of docking accuracy. \emph{Journal of Medicinal Chemistry} \textbf{47}, 1739--1749 (2004).

\bibitem{halgren2004glide} Halgren, T. A. et al. Glide: a new approach for rapid, accurate docking and scoring. 2. Enrichment factors in database screening. \emph{Journal of Medicinal Chemistry} \textbf{47}, 1750--1759 (2004).

\bibitem{jones1997development} Jones, G., Willett, P., Glen, R. C., Leach, A. R. \& Taylor, R. Development and validation of a genetic algorithm for flexible docking. \emph{Journal of Molecular Biology} \textbf{267}, 727--748 (1997).

\bibitem{rarey1996fast} Rarey, M., Kramer, B., Lengauer, T. \& Klebe, G. A fast flexible docking method using an incremental construction algorithm. \emph{Journal of Molecular Biology} \textbf{261}, 470--489 (1996).

\bibitem{goodsell1990automated} Goodsell, D. S. \& Olson, A. J. Automated docking of substrates to proteins by simulated annealing. \emph{Proteins: Structure, Function, and Bioinformatics} \textbf{8}, 195--202 (1990).

\bibitem{clark2006consensus} Clark, R. D., Strizhev, A., Leonard, J. M., Blake, J. F. \& Matthew, J. B. Consensus scoring for ligand/protein interactions. \emph{Journal of Molecular Graphics and Modelling} \textbf{20}, 281--295 (2002).

\bibitem{charifson1999consensus} Charifson, P. S., Corkery, J. J., Murcko, M. A. \& Walters, W. P. Consensus scoring: a method for obtaining improved hit rates from docking databases of three-dimensional structures into proteins. \emph{Journal of Medicinal Chemistry} \textbf{42}, 5100--5109 (1999).

\bibitem{wang2003further} Wang, R., Lu, Y. \& Wang, S. Comparative evaluation of 11 scoring functions for molecular docking. \emph{Journal of Medicinal Chemistry} \textbf{46}, 2287--2303 (2003).

\bibitem{eldridge1997empirical} Eldridge, M. D., Murray, C. W., Auton, T. R., Paolini, G. V. \& Mee, R. P. Empirical scoring functions: I. The development of a fast empirical scoring function to estimate the binding affinity of ligands in receptor complexes. \emph{Journal of Computer-Aided Molecular Design} \textbf{11}, 425--445 (1997).

\bibitem{bohm1994novel} Böhm, H.-J. The development of a simple empirical scoring function to estimate the binding constant for a protein-ligand complex of known three-dimensional structure. \emph{Journal of Computer-Aided Molecular Design} \textbf{8}, 243--256 (1994).

\bibitem{head1996validate} Head, R. D., Smythe, M. L., Oprea, T. I., Waller, C. L. \& Green, S. M. VALIDATE: a new method for the receptor-based prediction of binding affinities of novel ligands. \emph{Journal of the American Chemical Society} \textbf{118}, 3959--3969 (1996).

\bibitem{jain1996scoring} Jain, A. N. Scoring noncovalent protein-ligand interactions: a continuous differentiable function tuned to compute binding affinities. \emph{Journal of Computer-Aided Molecular Design} \textbf{10}, 427--440 (1996).

\bibitem{gehlhaar1995molecular} Gehlhaar, D. K. et al. Molecular recognition of the inhibitor AG-1343 by HIV-1 protease: conformationally flexible docking by evolutionary programming. \emph{Chemistry \& Biology} \textbf{2}, 317--324 (1995).

\bibitem{muegge1999pmf} Muegge, I. \& Martin, Y. C. A general and fast scoring function for protein-ligand interactions: a simplified potential approach. \emph{Journal of Medicinal Chemistry} \textbf{42}, 791--804 (1999).

\bibitem{mitchell1999bleep} Mitchell, J. B. O., Laskowski, R. A., Alex, A. \& Thornton, J. M. BLEEP--potential of mean force describing protein-ligand interactions: I. Generating potential. \emph{Journal of Computational Chemistry} \textbf{20}, 1165--1176 (1999).

\bibitem{gohlke2000knowledge} Gohlke, H., Hendlich, M. \& Klebe, G. Knowledge-based scoring function to predict protein-ligand interactions. \emph{Journal of Molecular Biology} \textbf{295}, 337--356 (2000).

\bibitem{velec2005drugscore} Velec, H. F. G., Gohlke, H. \& Klebe, G. DrugScore(CSD)-knowledge-based scoring function derived from small molecule crystal data with superior recognition rate of near-native ligand poses and better affinity prediction. \emph{Journal of Medicinal Chemistry} \textbf{48}, 6296--6303 (2005).

\bibitem{raha2000prediction} Raha, K. & Merz Jr, K. M. A quantum mechanics-based scoring function: study of zinc ion-mediated ligand binding. \emph{Journal of the American Chemical Society} \textbf{126}, 1020--1021 (2004).

\bibitem{ferrara2004assessing} Ferrara, P., Gohlke, H., Price, D. J., Klebe, G. \& Brooks III, C. L. Assessing scoring functions for protein-ligand interactions. \emph{Journal of Medicinal Chemistry} \textbf{47}, 3032--3047 (2004).

\bibitem{warren2006comparative} Warren, G. L. et al. A critical assessment of docking programs and scoring functions. \emph{Journal of Medicinal Chemistry} \textbf{49}, 5912--5931 (2006).

\bibitem{cross2009comparison} Cross, J. B. et al. Comparison of several molecular docking programs: pose prediction and virtual screening accuracy. \emph{Journal of Chemical Information and Modeling} \textbf{49}, 1455--1474 (2009).

\bibitem{cheng2009comparative} Cheng, T., Li, X., Li, Y., Liu, Z. \& Wang, R. Comparative assessment of scoring functions on a diverse test set. \emph{Journal of Chemical Information and Modeling} \textbf{49}, 1079--1093 (2009).

\bibitem{mysinger2012directory} Mysinger, M. M., Carchia, M., Irwin, J. J. \& Shoichet, B. K. Directory of useful decoys, enhanced (DUD-E): better ligands and decoys for better benchmarking. \emph{Journal of Medicinal Chemistry} \textbf{55}, 6582--6594 (2012).

\bibitem{rohrer2009maximum} Rohrer, S. G. \& Baumann, K. Maximum unbiased validation (MUV) data sets for virtual screening based on PubChem bioactivity data. \emph{Journal of Chemical Information and Modeling} \textbf{49}, 169--184 (2009).

\bibitem{lagerstrom2008structural} Lagerström, M. C. \& Schiöth, H. B. Structural diversity of G protein-coupled receptors and significance for drug discovery. \emph{Nature Reviews Drug Discovery} \textbf{7}, 339--357 (2008).

\bibitem{overington2006many} Overington, J. P., Al-Lazikani, B. \& Hopkins, A. L. How many drug targets are there? \emph{Nature Reviews Drug Discovery} \textbf{5}, 993--996 (2006).

\bibitem{rask2001genome} Rask-Andersen, M., Almén, M. S. \& Schiöth, H. B. Trends in the exploitation of novel drug targets. \emph{Nature Reviews Drug Discovery} \textbf{10}, 579--590 (2011).

\bibitem{hopkins2002druggable} Hopkins, A. L. \& Groom, C. R. The druggable genome. \emph{Nature Reviews Drug Discovery} \textbf{1}, 727--730 (2002).

\bibitem{imming2006drugs} Imming, P., Sinning, C. \& Meyer, A. Drugs, their targets and the nature and number of drug targets. \emph{Nature Reviews Drug Discovery} \textbf{5}, 821--834 (2006).

\bibitem{mestres2008computational} Mestres, J., Gregori-Puigjané, E., Valverde, S. \& Solé, R. V. Data completeness--the Achilles heel of drug-target networks. \emph{Nature Biotechnology} \textbf{26}, 983--984 (2008).

\bibitem{keiser2007relating} Keiser, M. J. et al. Relating protein pharmacology by ligand chemistry. \emph{Nature Biotechnology} \textbf{25}, 197--206 (2007).

\bibitem{keiser2009predicting} Keiser, M. J. et al. Predicting new molecular targets for known drugs. \emph{Nature} \textbf{462}, 175--181 (2009).

\bibitem{campillos2008drug} Campillos, M., Kuhn, M., Gavin, A.-C., Jensen, L. J. \& Bork, P. Drug target identification using side-effect similarity. \emph{Science} \textbf{321}, 263--266 (2008).

\bibitem{iorio2010discovery} Iorio, F. et al. Discovery of drug mode of action and drug repositioning from transcriptional responses. \emph{Proceedings of the National Academy of Sciences} \textbf{107}, 14621--14626 (2010).

\bibitem{lamb2006connectivity} Lamb, J. et al. The Connectivity Map: using gene-expression signatures to connect small molecules, genes, and disease. \emph{Science} \textbf{313}, 1929--1935 (2006).

\bibitem{subramanian2017next} Subramanian, A. et al. A next generation connectivity map: L1000 platform and the first 1,000,000 profiles. \emph{Cell} \textbf{171}, 1437--1452 (2017).

\bibitem{moffat2017opportunities} Moffat, J. G., Vincent, F., Lee, J. A., Eder, J. \& Prunotto, M. Opportunities and challenges in phenotypic drug discovery: an industry perspective. \emph{Nature Reviews Drug Discovery} \textbf{16}, 531--543 (2017).

\bibitem{swinney2011phenotypic} Swinney, D. C. \& Anthony, J. How were new medicines discovered? \emph{Nature Reviews Drug Discovery} \textbf{10}, 507--519 (2011).

\bibitem{eder2014application} Eder, J., Sedrani, R. \& Wiesmann, C. The discovery of first-in-class drugs: origins and evolution. \emph{Nature Reviews Drug Discovery} \textbf{13}, 577--587 (2014).

\bibitem{clark2014drug} Clark, D. E. What has computer-aided molecular design ever done for drug discovery? \emph{Expert Opinion on Drug Discovery} \textbf{1}, 103--110 (2006).

\bibitem{schneider2018automating} Schneider, G. Automating drug discovery. \emph{Nature Reviews Drug Discovery} \textbf{17}, 97--113 (2018).

\bibitem{bajorath2002integration} Bajorath, J. Integration of virtual and high-throughput screening. \emph{Nature Reviews Drug Discovery} \textbf{1}, 882--894 (2002).

\bibitem{klebe2006virtual} Klebe, G. Virtual ligand screening: strategies, perspectives and limitations. \emph{Drug Discovery Today} \textbf{11}, 580--594 (2006).

\bibitem{kolb2009matching} Kolb, P. et al. Structure-based discovery of β2-adrenergic receptor ligands. \emph{Proceedings of the National Academy of Sciences} \textbf{106}, 6843--6848 (2009).

\bibitem{carlsson2011ligand} Carlsson, J. et al. Ligand discovery from a dopamine D3 receptor homology model and crystal structure. \emph{Nature Chemical Biology} \textbf{7}, 769--778 (2011).

\bibitem{katritch2010analysis} Katritch, V., Rueda, M., Lam, P. C.-H., Yeager, M. \& Abagyan, R. GPCR 3D homology models for ligand screening: lessons learned from blind predictions of adenosine A2a receptor complex. \emph{Proteins: Structure, Function, and Bioinformatics} \textbf{78}, 197--211 (2010).

\bibitem{de2012structure} de Graaf, C. et al. Structure-based discovery of allosteric modulators of two related class B G-protein-coupled receptors. \emph{ChemMedChem} \textbf{6}, 2159--2169 (2011).

\bibitem{wang2013comprehensive} Wang, C. et al. Comprehensive evaluation of ten docking programs on a diverse set of protein-ligand complexes: the prediction accuracy of sampling power and scoring power. \emph{Physical Chemistry Chemical Physics} \textbf{18}, 12964--12975 (2016).

\bibitem{genheden2015g} Genheden, S. \& Ryde, U. The MM/PBSA and MM/GBSA methods to estimate ligand-binding affinities. \emph{Expert Opinion on Drug Discovery} \textbf{10}, 449--461 (2015).

\bibitem{sun2014assessing} Sun, H., Li, Y., Tian, S., Xu, L. \& Hou, T. Assessing the performance of MM/PBSA and MM/GBSA methods. 4. Accuracies of MM/PBSA and MM/GBSA methodologies evaluated by various simulation protocols using PDBbind data set. \emph{Physical Chemistry Chemical Physics} \textbf{16}, 16719--16729 (2014).

\bibitem{wang2015recent} Wang, E. et al. End-point binding free energy calculation with MM/PBSA and MM/GBSA: strategies and applications in drug design. \emph{Chemical Reviews} \textbf{119}, 9478--9508 (2019).

\bibitem{shirts2003equilibrium} Shirts, M. R. \& Pande, V. S. Comparison of efficiency and bias of free energies computed by exponential averaging, the Bennett acceptance ratio, and thermodynamic integration. \emph{Journal of Chemical Physics} \textbf{122}, 144107 (2005).

\bibitem{bennett1976efficient} Bennett, C. H. Efficient estimation of free energy differences from Monte Carlo data. \emph{Journal of Computational Physics} \textbf{22}, 245--268 (1976).

\bibitem{zwanzig1954high} Zwanzig, R. W. High‐temperature equation of state by a perturbation method. I. Nonpolar gases. \emph{Journal of Chemical Physics} \textbf{22}, 1420--1426 (1954).

\bibitem{kirkwood1935statistical} Kirkwood, J. G. Statistical mechanics of fluid mixtures. \emph{Journal of Chemical Physics} \textbf{3}, 300--313 (1935).

\bibitem{hansen2021theory} Hansen, J.-P. \& McDonald, I. R. \emph{Theory of simple liquids: with applications to soft matter}. Academic Press (2013).

\bibitem{cummins2005chemical} Cummins, D. J. \& Bell, M. A. Integrating everything: the molecule selection toolkit, a system for compound prioritization in drug discovery. \emph{Journal of Chemical Information and Modeling} \textbf{56}, 2077--2086 (2016).

\bibitem{jin2005evolutionary} Jin, Y. \& Branke, J. Evolutionary optimization in uncertain environments--a survey. \emph{IEEE Transactions on Evolutionary Computation} \textbf{9}, 303--317 (2005).

\bibitem{koza1992genetic} Koza, J. R. \emph{Genetic Programming: On the Programming of Computers by Means of Natural Selection}. MIT Press (1992).

\bibitem{cranmer2020discovering} Cranmer, M. et al. Discovering symbolic models from deep learning with inductive biases. \emph{Advances in Neural Information Processing Systems} \textbf{33}, 17429--17442 (2020).

\bibitem{swanson1986fish} Swanson, D. R. Fish oil, Raynaud's syndrome, and undiscovered public knowledge. \emph{Perspectives in Biology and Medicine} \textbf{30}, 7--18 (1986).

\bibitem{hastings2013chebi} Hastings, J. et al. The ChEBI reference database and ontology for biologically relevant chemistry: enhancements for 2013. \emph{Nucleic Acids Research} \textbf{41}, D456--D463 (2013).
\end{thebibliography}

\end{document}
