## Supplementary material for "AMODO-EO: A Practical Framework for Adaptive Objective Discovery in Multi-Objective Drug Optimization": Source code of the algorithm: minimal_test.py

```
"""
Minimal test for objectives issue
"""

print("Testing objectives handling...")

# Test the scoring method directly
import pandas as pd
import numpy as np

# Mock data
df = pd.DataFrame({
    'feature_0': [1, 2, 3],
    'feature_1': [4, 5, 6],
    'binding_affinity': [0.5, 0.6, 0.7],
    'admet_score': [0.4, 0.5, 0.6],
    'drug_likeness': [0.3, 0.4, 0.5]
})

# Test with list of objectives
objectives_list = ['objective_0', 'objective_1', 'objective_2']

print(f"DataFrame shape: {df.shape}")
print(f"Objectives: {objectives_list}")
print(f"Objectives type: {type(objectives_list)}")

# Test if-else logic
if isinstance(objectives_list, dict):
    print("Is dict")
elif isinstance(objectives_list, list):
    print("Is list - this should print")
else:
    print(f"Unknown type: {type(objectives_list)}")

print("✅ Basic test completed")
```
