## Supplementary material for "AMODO-EO: A Practical Framework for Adaptive Objective Discovery in Multi-Objective Drug Optimization": Source code of the algorithm: quick_test.py

```
"""
Quick validation test for the objectives issue
"""

import sys
sys.path.append('src')

import pandas as pd
import numpy as np
from src.real_datasets import RealDatasetManager
from src.publication_ready_amodo_heo import PublicationReadyAMODOHEO

try:
    print("Testing the objectives fix...")
    
    # Create test data
    dataset_manager = RealDatasetManager()
    X, y = dataset_manager.get_dataset('benchmark', n_compounds=20)
    
    # Create DataFrame
    df = pd.DataFrame(X, columns=[f'feature_{i}' for i in range(X.shape[1])])
    objectives = [f'objective_{i}' for i in range(y.shape[1])]
    
    # Initialize AMODO-HEO
    amodo = PublicationReadyAMODOHEO(
        max_objectives=3,
        significance_threshold=0.1,
        min_improvement=0.01
    )
    
    print(f"Created dataset: {df.shape}")
    print(f"Objectives: {objectives}")
    
    # Test the scoring function directly
    scores = amodo._score_compounds_with_objectives(df, objectives)
    print(f"✅ Scoring works: {scores.shape}")
    
    # Test full optimization
    result = amodo.optimize_compounds(df, objectives)
    print(f"✅ Full optimization works!")
    
except Exception as e:
    print(f"❌ Error: {e}")
    import traceback
    traceback.print_exc()
```
