## Supplementary material for "AMODO-EO: A Practical Framework for Adaptive Objective Discovery in Multi-Objective Drug Optimization": Source code of the algorithm: README.md

# AMODO-EO: Adaptive Multi-Objective Drug Optimization with Emergent Objectives

**Authors:** Shetty Nihal Naveen, B. Karthik Udupa, Ashil A. Shetty  
**Institution:** Department of Information Science & Engineering, Sahyadri College of Engineering and Management, Mangaluru, Karnataka, India

## 📄 Paper

This repository contains the complete source code and manuscript for our paper:

**"AMODO-EO: A Practical Framework for Adaptive Objective Discovery in Multi-Objective Drug Optimization"**

## 🗂️ Repository Structure

```
├── manuscript/
│   ├── amodo_heo_comprehensive_research_paper.tex  # Main manuscript
│   ├── references.bib                              # Bibliography
│   └── supplementary_materials.tex                 # Supplementary materials
├── src/
│   ├── publication_ready_amodo_heo.py             # Main AMODO-EO algorithm
│   ├── statistical_benchmarking.py               # Benchmarking framework
│   ├── real_datasets.py                          # Dataset management
│   └── comprehensive_benchmark.py                # Complete benchmark suite
├── experiments/
│   └── experimental_validation.py                # Validation framework
├── results/
│   ├── final_objectives.json                     # Discovered objectives
│   └── test_results.json                         # Experimental results
├── requirements.txt                               # Python dependencies
└── README.md                                      # This file
```

## 🚀 Quick Start

### Installation
```bash
pip install -r requirements.txt
```

### Run AMODO-EO
```python
from src.publication_ready_amodo_heo import PublicationReadyAMODOHEO

# Initialize algorithm
amodo = PublicationReadyAMODOHEO(
    population_size=100,
    generations=500,
    discovery_enabled=True
)

# Run optimization with objective discovery
results = amodo.optimize(X, y)
print(f"Discovered objectives: {results['discovered_objectives']}")
```

### Run Benchmarks
```python
from src.comprehensive_benchmark import ComprehensiveBenchmark

benchmark = ComprehensiveBenchmark()
results = benchmark.run_full_benchmark()
```

### Reproduce Paper Results
```bash
python experiments/experimental_validation.py
python run_publication_benchmark.py
python test_publication_readiness.py
```

## 📊 Key Results

- **Discovered 6 meaningful objectives** across datasets
- **HBA/RTB ratio** found in 100% of runs (highly robust)
- **2.9% performance cost** for discovery capability
- **Statistical validation** confirms objective independence

## 📜 Citation

```bibtex
@article{shetty2025amodo,
  title={AMODO-EO: A Practical Framework for Adaptive Objective Discovery in Multi-Objective Drug Optimization},
  author={Shetty, Nihal Naveen and Udupa, B. Karthik and Shetty, Ashil A.},
  journal={arXiv preprint arXiv:2408.xxxxx},
  year={2025}
}
```

## 📧 Contact

- **Shetty Nihal Naveen:**
- **B. Karthik Udupa:**  
- **Ashil A. Shetty:**

## 🏫 Affiliation

Department of Information Science & Engineering  
Sahyadri College of Engineering and Management  
Mangaluru, Karnataka, India

## 📄 License

This project is available under the MIT License. See LICENSE file for details.
