## Supplementary material for "AMODO-EO: A Practical Framework for Adaptive Objective Discovery in Multi-Objective Drug Optimization": Source code of the algorithm: run_publication_benchmark.py

```
"""
Publication-Ready Comprehensive Benchmark Runner
==============================================

This script runs the complete benchmarking suite for AMODO-HEO publication,
including all modern baselines, real datasets, and statistical analysis.

Author: Nihal Shetty
Date: August 4, 2025
"""

import sys
import os
sys.path.append(os.path.dirname(os.path.abspath(__file__)))

import numpy as np
import pandas as pd
import time
from pathlib import Path

# Import our modules
from src.real_datasets import RealDatasetManager
from src.modern_baselines import get_all_baseline_optimizers
from src.publication_ready_amodo_heo import PublicationReadyAMODOHEO
from src.comprehensive_benchmark import ComprehensiveBenchmark

def run_publication_benchmark():
    """
    Run the complete publication-ready benchmark
    """
    
    print("="*80)
    print("AMODO-HEO PUBLICATION-READY COMPREHENSIVE BENCHMARK")
    print("="*80)
    print("Version: 2.0")
    print("Date: August 4, 2025")
    print("Objective: Generate publication-ready results for top-tier journals")
    print("="*80)
    
    # Initialize components
    benchmark = ComprehensiveBenchmark(results_dir="results/publication_benchmark")
    
    # Define comprehensive benchmark parameters
    benchmark_config = {
        'n_folds': 5,  # 5-fold cross-validation
        'datasets': [
            'benchmark',      # 10,000 compounds from multiple sources
            'chembl_small',   # 2,000 ChEMBL compounds
            'chembl_large',   # 5,000 ChEMBL compounds
        ],
        'algorithms': [
            'amodo_heo',      # Our algorithm
            'nsga2',          # Gold standard MOO
            'moead',          # Modern decomposition-based MOO
            'random_forest',  # ML baseline
            'xgboost',        # Modern ML baseline
            'random',         # Simple baseline
            'weighted_sum',   # Traditional baseline
        ]
    }
    
    print(f"Benchmark Configuration:")
    print(f"  Cross-validation folds: {benchmark_config['n_folds']}")
    print(f"  Datasets: {benchmark_config['datasets']}")
    print(f"  Algorithms: {benchmark_config['algorithms']}")
    print(f"  Total experiments: {len(benchmark_config['datasets']) * len(benchmark_config['algorithms']) * benchmark_config['n_folds']}")
    
    # Estimate runtime
    estimated_time = len(benchmark_config['datasets']) * len(benchmark_config['algorithms']) * benchmark_config['n_folds'] * 30  # 30 seconds per experiment
    print(f"  Estimated runtime: {estimated_time // 60:.0f} minutes")
    
    print("\n" + "="*80)
    print("STARTING COMPREHENSIVE BENCHMARK")
    print("="*80)
    
    start_time = time.time()
    
    # Run the benchmark
    results = benchmark.run_comprehensive_benchmark(
        n_folds=benchmark_config['n_folds'],
        datasets=benchmark_config['datasets'],
        algorithms=benchmark_config['algorithms']
    )
    
    total_time = time.time() - start_time
    
    print("\n" + "="*80)
    print("BENCHMARK COMPLETED")
    print("="*80)
    print(f"Total runtime: {total_time // 60:.0f} minutes {total_time % 60:.0f} seconds")
    
    # Extract key results
    publication_readiness = results['publication_ready']
    summary = results['summary_report']
    
    print("\n" + "="*60)
    print("PUBLICATION READINESS ASSESSMENT")
    print("="*60)
    
    print(f"Overall Score: {publication_readiness['overall_score']:.1f}/100")
    print(f"Publication Ready: {'✅ YES' if publication_readiness['publication_ready'] else '❌ NO'}")
    
    print(f"\nCriteria Breakdown:")
    for criterion, details in publication_readiness['criteria'].items():
        print(f"  {criterion}: {details['score']:.1f}/{details['max_score']} - {details['note']}")
    
    print(f"\nRecommendations:")
    for rec in publication_readiness['recommendations']:
        print(f"  • {rec}")
    
    print("\n" + "="*60)
    print("PERFORMANCE COMPARISON")
    print("="*60)
    
    # Overall algorithm ranking
    ranking = summary['performance_comparison']['overall_ranking']
    print(f"Algorithm Rankings:")
    for i, alg_perf in enumerate(ranking):
        marker = "🏆" if i == 0 else "🥈" if i == 1 else "🥉" if i == 2 else "  "
        print(f"  {marker} {i+1}. {alg_perf['algorithm']}: {alg_perf['mean_performance']:.4f} ± {alg_perf['std_performance']:.4f}")
    
    # AMODO-HEO specific results
    amodo_rank = summary['performance_comparison']['amodo_heo_rank']
    if amodo_rank:
        print(f"\nAMODO-HEO Performance:")
        print(f"  Rank: #{amodo_rank} out of {len(ranking)} algorithms")
        
        # Find AMODO-HEO details
        amodo_details = next((alg for alg in ranking if alg['algorithm'] == 'amodo_heo'), None)
        if amodo_details:
            print(f"  Mean Performance: {amodo_details['mean_performance']:.4f} ± {amodo_details['std_performance']:.4f}")
            print(f"  Execution Time: {amodo_details['mean_execution_time']:.2f}s")
            print(f"  Datasets Tested: {amodo_details['datasets_tested']}")
    
    print("\n" + "="*60)
    print("STATISTICAL SIGNIFICANCE")
    print("="*60)
    
    stat_sig = summary['statistical_significance']
    print(f"Significant Improvements: {stat_sig['significant_improvements']}/{stat_sig['total_comparisons']}")
    print(f"Improvement Rate: {stat_sig['improvement_rate']:.1%}")
    
    # Detailed statistical results
    statistical_results = results['statistical_analysis']
    
    print(f"\nDetailed Statistical Results:")
    for dataset_name, pairwise_results in statistical_results['pairwise_comparisons'].items():
        print(f"\n  Dataset: {dataset_name}")
        
        amodo_comparisons = {k: v for k, v in pairwise_results.items() if 'amodo_heo' in k}
        
        if amodo_comparisons:
            print(f"    AMODO-HEO Comparisons:")
            for comparison, result in amodo_comparisons.items():
                algorithms = comparison.split('_vs_')
                if algorithms[0] == 'amodo_heo':
                    direction = "beats" if result['mean_diff'] > 0 else "loses to"
                    other_alg = algorithms[1]
                else:
                    direction = "beats" if result['mean_diff'] < 0 else "loses to"
                    other_alg = algorithms[0]
                
                significance = "✅" if result['significant'] else "❌"
                effect_size_label = "Large" if abs(result['effect_size']) > 0.8 else "Medium" if abs(result['effect_size']) > 0.5 else "Small"
                
                print(f"      {significance} AMODO-HEO {direction} {other_alg}")
                print(f"         p={result['p_value']:.4f}, Cohen's d={result['effect_size']:.3f} ({effect_size_label})")
    
    print("\n" + "="*60)
    print("PUBLICATION RECOMMENDATIONS")
    print("="*60)
    
    if publication_readiness['overall_score'] >= 85:
        print("🎉 EXCELLENT! Ready for top-tier publication (Nature, Science, Cell)")
        target_journals = [
            "Nature Machine Intelligence",
            "Nature Methods",
            "Science Advances",
            "Cell Systems"
        ]
    elif publication_readiness['overall_score'] >= 75:
        print("✅ GOOD! Ready for high-impact specialized journals")
        target_journals = [
            "Journal of Chemical Information and Modeling",
            "Bioinformatics",
            "Journal of Medicinal Chemistry",
            "Drug Discovery Today"
        ]
    elif publication_readiness['overall_score'] >= 60:
        print("⚠️ NEEDS IMPROVEMENTS for top journals, suitable for specialized venues")
        target_journals = [
            "Computational Biology and Chemistry",
            "Current Computer-Aided Drug Design",
            "Molecular Informatics"
        ]
    else:
        print("❌ NEEDS MAJOR IMPROVEMENTS before publication")
        target_journals = [
            "Consider as technical report first",
            "Submit to workshops/conferences for feedback"
        ]
    
    print(f"\nRecommended Target Journals:")
    for journal in target_journals:
        print(f"  • {journal}")
    
    print("\n" + "="*60)
    print("NEXT STEPS FOR PUBLICATION")
    print("="*60)
    
    if publication_readiness['overall_score'] >= 70:
        print("✅ Ready to proceed with manuscript preparation!")
        print("\nManuscript Structure Recommendations:")
        print("  1. Abstract: Emphasize statistical significance and performance gains")
        print("  2. Introduction: Position as advancement over NSGA-II/MOEA-D")
        print("  3. Methods: Detail objective discovery and validation methodology")
        print("  4. Results: Present cross-validation results and statistical tests")
        print("  5. Discussion: Compare with state-of-the-art, acknowledge limitations")
        print("  6. Conclusion: Emphasize practical impact for drug discovery")
        
        print(f"\nKey Numbers for Abstract:")
        if amodo_details:
            improvement_vs_baseline = ((amodo_details['mean_performance'] - 0.5) / 0.5) * 100
            print(f"  • Performance improvement: {improvement_vs_baseline:.1f}% over baseline")
        print(f"  • Statistical significance: {stat_sig['improvement_rate']:.0%} of comparisons significant")
        print(f"  • Cross-validation: {benchmark_config['n_folds']}-fold CV across {len(benchmark_config['datasets'])} datasets")
        print(f"  • Benchmarks: Compared against {len(benchmark_config['algorithms'])-1} state-of-the-art algorithms")
    
    else:
        print("⚠️ Improvements needed before publication:")
        
        # Specific recommendations based on criteria scores
        criteria = publication_readiness['criteria']
        
        if criteria.get('statistical_significance', {}).get('score', 0) < 20:
            print("  • Improve statistical significance (add more datasets or refine algorithm)")
        
        if criteria.get('modern_baselines', {}).get('score', 0) < 20:
            print("  • Add more modern baseline comparisons")
        
        if criteria.get('dataset_diversity', {}).get('score', 0) < 15:
            print("  • Test on more diverse datasets")
        
        if criteria.get('effect_sizes', {}).get('score', 0) < 7:
            print("  • Work on increasing effect sizes (more substantial improvements)")
    
    print("\n" + "="*80)
    print("BENCHMARK COMPLETE - RESULTS SAVED")
    print("="*80)
    print(f"Results directory: {benchmark.results_dir}")
    print("Files generated:")
    print("  • benchmark_results_[timestamp].json - Complete raw results")
    print("  • publication_summary_[timestamp].txt - Publication-ready summary")
    
    return results

if __name__ == "__main__":
    # Check if required packages are available
    try:
        import sklearn
        import scipy
        import numpy
        import pandas
        print("✅ All required packages available")
    except ImportError as e:
        print(f"❌ Missing required package: {e}")
        print("Please install: pip install scikit-learn scipy numpy pandas")
        sys.exit(1)
    
    # Run the complete benchmark
    results = run_publication_benchmark()
    
    # Final summary
    print(f"\n🎯 FINAL VERDICT:")
    if results['publication_ready']['publication_ready']:
        print("🎉 AMODO-HEO IS READY FOR PUBLICATION!")
    else:
        print("⚠️  AMODO-HEO needs improvements for top-tier publication")
    
    print(f"📊 Overall Score: {results['publication_ready']['overall_score']:.1f}/100")
```
