## Supplementary material for "AMODO-EO: A Practical Framework for Adaptive Objective Discovery in Multi-Objective Drug Optimization": Source code of the algorithm: setup_publication.py

```
"""
AMODO-HEO Publication Setup
===========================

Setup script to prepare the publication-ready AMODO-HEO system.

Author: Nihal Shetty
Date: August 4, 2025
"""

import os
import sys
import subprocess
import json
from pathlib import Path

def create_directory_structure():
    """Create necessary directory structure"""
    print("Creating directory structure...")
    
    try:
        directories = [
            "src",
            "results",
            "results/publication_benchmark",
            "results/benchmarks",
            "results/figures",
            "results/reports",
            "data",
            "data/temp",
            "data/cache",
            "logs"
        ]
        
        for directory in directories:
            os.makedirs(directory, exist_ok=True)
            print(f"✅ Created directory: {directory}")
        
        return True
        
    except Exception as e:
        print(f"❌ Error creating directories: {e}")
        return False

def check_python_version():
    """Check Python version compatibility"""
    print("Checking Python version...")
    
    version = sys.version_info
    if version.major == 3 and version.minor >= 8:
        print(f"✅ Python {version.major}.{version.minor}.{version.micro} - Compatible")
        return True
    else:
        print(f"❌ Python {version.major}.{version.minor}.{version.micro} - Requires Python 3.8+")
        return False

def install_dependencies():
    """Install required dependencies"""
    print("Installing dependencies...")
    
    try:
        # Check if requirements.txt exists
        if not os.path.exists("requirements.txt"):
            print("❌ requirements.txt not found")
            return False
        
        # Install dependencies
        result = subprocess.run([
            sys.executable, "-m", "pip", "install", "-r", "requirements.txt"
        ], capture_output=True, text=True)
        
        if result.returncode == 0:
            print("✅ Dependencies installed successfully")
            return True
        else:
            print(f"❌ Failed to install dependencies: {result.stderr}")
            return False
            
    except Exception as e:
        print(f"❌ Error installing dependencies: {e}")
        return False

def verify_core_files():
    """Verify that all core files are present"""
    print("Verifying core files...")
    
    required_files = [
        "src/real_datasets.py",
        "src/modern_baselines.py", 
        "src/comprehensive_benchmark.py",
        "src/publication_ready_amodo_heo.py",
        "run_publication_benchmark.py",
        "test_publication_readiness.py",
        "requirements.txt"
    ]
    
    missing_files = []
    
    for file_path in required_files:
        if os.path.exists(file_path):
            print(f"✅ Found: {file_path}")
        else:
            print(f"❌ Missing: {file_path}")
            missing_files.append(file_path)
    
    if missing_files:
        print(f"\n❌ Missing {len(missing_files)} required files")
        return False
    else:
        print(f"\n✅ All {len(required_files)} required files present")
        return True

def create_configuration():
    """Create default configuration file"""
    print("Creating configuration...")
    
    try:
        config = {
            "amodo_heo": {
                "population_size": 100,
                "n_generations": 50,
                "mutation_rate": 0.1,
                "crossover_rate": 0.8,
                "objective_discovery_rate": 0.2
            },
            "benchmark": {
                "n_folds": 5,
                "test_size": 0.2,
                "random_state": 42,
                "n_jobs": -1
            },
            "datasets": {
                "benchmark_size": 10000,
                "chembl_small_size": 2000,
                "chembl_large_size": 5000,
                "cache_enabled": True
            },
            "output": {
                "results_dir": "results",
                "save_figures": True,
                "save_raw_data": True,
                "publication_format": True
            }
        }
        
        os.makedirs("config", exist_ok=True)
        
        with open("config/config.json", "w") as f:
            json.dump(config, f, indent=2)
        
        print("✅ Configuration created: config/config.json")
        return True
        
    except Exception as e:
        print(f"❌ Error creating configuration: {e}")
        return False

def run_setup():
    """Run complete setup process"""
    print("="*80)
    print("AMODO-HEO PUBLICATION SETUP")
    print("="*80)
    print("Version: 2.0")
    print("Date: August 4, 2025")
    print("="*80)
    
    setup_steps = [
        ("Python Version Check", check_python_version),
        ("Directory Structure", create_directory_structure),
        ("Core Files Verification", verify_core_files),
        ("Configuration Creation", create_configuration),
        # ("Dependencies Installation", install_dependencies),  # Skip auto-install for now
    ]
    
    results = {}
    
    for step_name, step_func in setup_steps:
        print(f"\n{'='*20} {step_name} {'='*20}")
        
        try:
            result = step_func()
            results[step_name] = result
            
            if result:
                print(f"✅ {step_name}: SUCCESS")
            else:
                print(f"❌ {step_name}: FAILED")
                
        except Exception as e:
            print(f"❌ {step_name}: ERROR - {e}")
            results[step_name] = False
    
    # Summary
    print("\n" + "="*80)
    print("SETUP SUMMARY")
    print("="*80)
    
    passed_steps = sum(results.values())
    total_steps = len(results)
    success_rate = (passed_steps / total_steps) * 100
    
    print(f"Setup Steps: {passed_steps}/{total_steps} ({success_rate:.1f}%)")
    
    if success_rate >= 100:
        print("🎉 SETUP COMPLETE! System ready for testing and benchmarking")
        
        print("\n📋 NEXT STEPS:")
        print("1. Install dependencies manually if needed:")
        print("   pip install -r requirements.txt")
        print("\n2. Run quick test to verify everything works:")
        print("   python test_publication_readiness.py")
        print("\n3. If tests pass, run full publication benchmark:")
        print("   python run_publication_benchmark.py")
        print("\n4. Review results in results/publication_benchmark/")
        
    elif success_rate >= 80:
        print("⚠️  SETUP MOSTLY COMPLETE - Some issues detected")
        
        failed_steps = [step for step, result in results.items() if not result]
        print(f"\nFailed steps: {', '.join(failed_steps)}")
        
    else:
        print("❌ SETUP FAILED - Major issues detected")
        
        print("\nFailed steps:")
        for step, result in results.items():
            if not result:
                print(f"  • {step}")
    
    # Save setup results
    setup_log = {
        "timestamp": os.popen('echo %date% %time%').read().strip(),
        "setup_results": results,
        "success_rate": success_rate,
        "python_version": f"{sys.version_info.major}.{sys.version_info.minor}.{sys.version_info.micro}",
        "working_directory": os.getcwd()
    }
    
    os.makedirs("logs", exist_ok=True)
    with open("logs/setup_log.json", "w") as f:
        json.dump(setup_log, f, indent=2)
    
    print(f"\n📄 Setup log saved to: logs/setup_log.json")
    
    return results

if __name__ == "__main__":
    results = run_setup()
    
    # Final message
    if sum(results.values()) == len(results):
        print("\n🚀 READY TO PROCEED!")
        print("The AMODO-HEO publication system is now set up and ready for testing.")
    else:
        print("\n🔧 SETUP NEEDS ATTENTION")
        print("Please resolve the issues above before proceeding.")
```
