## Supplementary material for "AMODO-EO: A Practical Framework for Adaptive Objective Discovery in Multi-Objective Drug Optimization": Source code of the algorithm: modern_baselines.py

```
"""
Modern Multi-Objective Optimization Baselines
============================================

Implementation of state-of-the-art MOO algorithms for fair comparison:
- NSGA-II (Non-dominated Sorting Genetic Algorithm II)
- MOEA/D (Multi-Objective Evolutionary Algorithm based on Decomposition)
- SPEA2 (Strength Pareto Evolutionary Algorithm 2)
- SMS-EMOA (S-Metric Selection Evolutionary Multi-Objective Algorithm)
- Modern ML baselines (DeepChem-inspired, Random Forest, XGBoost)

Author: Nihal Shetty
Date: August 4, 2025
"""

import numpy as np
import pandas as pd
from typing import Dict, List, Tuple, Callable, Optional
from abc import ABC, abstractmethod
from sklearn.ensemble import RandomForestRegressor
from sklearn.multioutput import MultiOutputRegressor
from sklearn.model_selection import cross_val_score
import xgboost as xgb
from scipy.spatial.distance import cdist
from scipy.stats import rankdata
import time

class BaseOptimizer(ABC):
    """Base class for multi-objective optimizers"""
    
    def __init__(self, name: str):
        self.name = name
        
    @abstractmethod
    def optimize(self, compounds_df: pd.DataFrame, objectives: Dict[str, Callable],
                max_iterations: int = 100) -> Tuple[pd.DataFrame, Dict]:
        """Optimize compounds using the algorithm"""
        pass

class NSGAIIOptimizer(BaseOptimizer):
    """
    NSGA-II: Non-dominated Sorting Genetic Algorithm II
    
    The gold standard for multi-objective optimization.
    Reference: Deb et al. (2002) "A fast and elitist multiobjective genetic algorithm: NSGA-II"
    """
    
    def __init__(self, population_size: int = 100, mutation_rate: float = 0.1,
                 crossover_rate: float = 0.9):
        super().__init__("NSGA-II")
        self.population_size = population_size
        self.mutation_rate = mutation_rate
        self.crossover_rate = crossover_rate
    
    def optimize(self, compounds_df: pd.DataFrame, objectives: Dict[str, Callable],
                max_iterations: int = 100) -> Tuple[pd.DataFrame, Dict]:
        """
        Run NSGA-II optimization
        """
        start_time = time.time()
        
        # Calculate objective values for all compounds
        objective_values = self._calculate_objectives(compounds_df, objectives)
        
        # Perform non-dominated sorting
        fronts = self._non_dominated_sort(objective_values)
        
        # Calculate crowding distance
        crowding_distances = self._calculate_crowding_distance(objective_values, fronts)
        
        # Assign fitness based on Pareto rank and crowding distance
        fitness_scores = self._assign_fitness(fronts, crowding_distances)
        
        # Create scored dataframe
        scored_df = compounds_df.copy()
        scored_df['nsga2_score'] = fitness_scores
        scored_df['pareto_rank'] = self._get_pareto_ranks(fronts, len(compounds_df))
        scored_df['crowding_distance'] = crowding_distances
        
        # Sort by fitness (higher is better)
        scored_df = scored_df.sort_values('nsga2_score', ascending=False)
        
        execution_time = time.time() - start_time
        
        metrics = {
            'algorithm': 'NSGA-II',
            'execution_time': execution_time,
            'num_fronts': len(fronts),
            'pareto_optimal_solutions': len(fronts[0]) if fronts else 0,
            'hypervolume': self._calculate_hypervolume(objective_values, fronts[0] if fronts else []),
            'convergence_metric': self._calculate_convergence_metric(objective_values, fronts),
            'diversity_metric': np.mean(crowding_distances) if crowding_distances else 0
        }
        
        return scored_df, metrics
    
    def _calculate_objectives(self, compounds_df: pd.DataFrame, 
                            objectives: Dict[str, Callable]) -> np.ndarray:
        """Calculate objective values for all compounds"""
        objective_matrix = []
        
        for idx, row in compounds_df.iterrows():
            compound_dict = row.to_dict()
            obj_values = []
            
            for obj_func in objectives.values():
                try:
                    value = obj_func(compound_dict)
                    obj_values.append(value if isinstance(value, (int, float)) else 0.0)
                except:
                    obj_values.append(0.0)
            
            objective_matrix.append(obj_values)
        
        return np.array(objective_matrix)
    
    def _non_dominated_sort(self, objective_values: np.ndarray) -> List[List[int]]:
        """Perform non-dominated sorting (Fast Non-dominated Sort)"""
        n = len(objective_values)
        fronts = []
        
        # For each solution
        domination_count = np.zeros(n)  # Number of solutions that dominate this solution
        dominated_solutions = [[] for _ in range(n)]  # Solutions dominated by this solution
        
        # Calculate domination relationships
        for i in range(n):
            for j in range(n):
                if i != j:
                    if self._dominates(objective_values[i], objective_values[j]):
                        dominated_solutions[i].append(j)
                    elif self._dominates(objective_values[j], objective_values[i]):
                        domination_count[i] += 1
        
        # Find first front
        current_front = []
        for i in range(n):
            if domination_count[i] == 0:
                current_front.append(i)
        
        # Generate subsequent fronts
        while current_front:
            fronts.append(current_front[:])
            next_front = []
            
            for i in current_front:
                for j in dominated_solutions[i]:
                    domination_count[j] -= 1
                    if domination_count[j] == 0:
                        next_front.append(j)
            
            current_front = next_front
        
        return fronts
    
    def _dominates(self, solution_a: np.ndarray, solution_b: np.ndarray) -> bool:
        """Check if solution_a dominates solution_b (assuming maximization)"""
        return (np.all(solution_a >= solution_b) and np.any(solution_a > solution_b))
    
    def _calculate_crowding_distance(self, objective_values: np.ndarray, 
                                   fronts: List[List[int]]) -> np.ndarray:
        """Calculate crowding distance for diversity preservation"""
        n = len(objective_values)
        distances = np.zeros(n)
        
        for front in fronts:
            if len(front) <= 2:
                # Boundary solutions get infinite distance
                for idx in front:
                    distances[idx] = float('inf')
                continue
            
            front_objectives = objective_values[front]
            n_objectives = front_objectives.shape[1]
            
            # Calculate distance for each objective
            for m in range(n_objectives):
                # Sort by objective m
                sorted_indices = np.argsort(front_objectives[:, m])
                
                # Boundary points get infinite distance
                distances[front[sorted_indices[0]]] = float('inf')
                distances[front[sorted_indices[-1]]] = float('inf')
                
                # Calculate range
                obj_range = front_objectives[sorted_indices[-1], m] - front_objectives[sorted_indices[0], m]
                
                if obj_range == 0:
                    continue
                
                # Calculate crowding distance for intermediate points
                for i in range(1, len(sorted_indices) - 1):
                    if distances[front[sorted_indices[i]]] != float('inf'):
                        distance_contribution = (
                            front_objectives[sorted_indices[i + 1], m] - 
                            front_objectives[sorted_indices[i - 1], m]
                        ) / obj_range
                        distances[front[sorted_indices[i]]] += distance_contribution
        
        return distances
    
    def _assign_fitness(self, fronts: List[List[int]], 
                       crowding_distances: np.ndarray) -> np.ndarray:
        """Assign fitness based on Pareto rank and crowding distance"""
        n = sum(len(front) for front in fronts)
        fitness = np.zeros(n)
        
        for rank, front in enumerate(fronts):
            for idx in front:
                # Higher rank is worse, so we use negative rank
                # Higher crowding distance is better (promotes diversity)
                fitness[idx] = -rank + crowding_distances[idx] / 1000.0  # Scale crowding distance
        
        return fitness
    
    def _get_pareto_ranks(self, fronts: List[List[int]], n: int) -> np.ndarray:
        """Get Pareto rank for each solution"""
        ranks = np.zeros(n)
        for rank, front in enumerate(fronts):
            for idx in front:
                ranks[idx] = rank + 1  # 1-indexed ranks
        return ranks
    
    def _calculate_hypervolume(self, objective_values: np.ndarray, 
                             pareto_front: List[int]) -> float:
        """Calculate hypervolume indicator (simplified 2D version)"""
        if not pareto_front or objective_values.shape[1] != 2:
            return 0.0
        
        # Reference point (origin for maximization)
        ref_point = np.zeros(objective_values.shape[1])
        
        # Get Pareto front points
        pareto_points = objective_values[pareto_front]
        
        # Sort by first objective
        sorted_indices = np.argsort(pareto_points[:, 0])
        sorted_points = pareto_points[sorted_indices]
        
        # Calculate hypervolume
        hv = 0.0
        prev_x = ref_point[0]
        
        for point in sorted_points:
            width = point[0] - prev_x
            height = point[1] - ref_point[1]
            hv += width * max(0, height)
            prev_x = point[0]
        
        return hv
    
    def _calculate_convergence_metric(self, objective_values: np.ndarray,
                                    fronts: List[List[int]]) -> float:
        """Calculate convergence metric (distance to Pareto front)"""
        if not fronts:
            return float('inf')
        
        pareto_front = objective_values[fronts[0]]
        
        # Calculate average distance from all solutions to Pareto front
        distances = []
        for obj_val in objective_values:
            min_dist = np.min(np.linalg.norm(pareto_front - obj_val, axis=1))
            distances.append(min_dist)
        
        return np.mean(distances)

class MOEADOptimizer(BaseOptimizer):
    """
    MOEA/D: Multi-Objective Evolutionary Algorithm based on Decomposition
    
    Reference: Zhang & Li (2007) "MOEA/D: A Multiobjective Evolutionary Algorithm Based on Decomposition"
    """
    
    def __init__(self, n_neighbors: int = 10, weight_vectors: Optional[np.ndarray] = None):
        super().__init__("MOEA/D")
        self.n_neighbors = n_neighbors
        self.weight_vectors = weight_vectors
    
    def optimize(self, compounds_df: pd.DataFrame, objectives: Dict[str, Callable],
                max_iterations: int = 100) -> Tuple[pd.DataFrame, Dict]:
        """Run MOEA/D optimization"""
        start_time = time.time()
        
        # Generate weight vectors if not provided
        n_objectives = len(objectives)
        if self.weight_vectors is None:
            self.weight_vectors = self._generate_weight_vectors(n_objectives, len(compounds_df))
        
        # Calculate objective values
        objective_values = self._calculate_objectives(compounds_df, objectives)
        
        # Decompose problem using weighted sum approach
        decomposed_scores = self._decompose_objectives(objective_values, self.weight_vectors)
        
        # Select best solution for each subproblem
        best_indices = np.argmax(decomposed_scores, axis=1)
        
        # Create scored dataframe
        scored_df = compounds_df.copy()
        scored_df['moead_score'] = np.max(decomposed_scores, axis=0)
        scored_df['subproblem_rank'] = rankdata(-scored_df['moead_score'], method='ordinal')
        
        scored_df = scored_df.sort_values('moead_score', ascending=False)
        
        execution_time = time.time() - start_time
        
        metrics = {
            'algorithm': 'MOEA/D',
            'execution_time': execution_time,
            'n_subproblems': len(self.weight_vectors),
            'n_neighbors': self.n_neighbors,
            'decomposition_quality': self._evaluate_decomposition_quality(decomposed_scores),
            'diversity_metric': self._calculate_weight_diversity()
        }
        
        return scored_df, metrics
    
    def _calculate_objectives(self, compounds_df: pd.DataFrame, 
                            objectives: Dict[str, Callable]) -> np.ndarray:
        """Calculate objective values (same as NSGA-II)"""
        objective_matrix = []
        
        for idx, row in compounds_df.iterrows():
            compound_dict = row.to_dict()
            obj_values = []
            
            for obj_func in objectives.values():
                try:
                    value = obj_func(compound_dict)
                    obj_values.append(value if isinstance(value, (int, float)) else 0.0)
                except:
                    obj_values.append(0.0)
            
            objective_matrix.append(obj_values)
        
        return np.array(objective_matrix)
    
    def _generate_weight_vectors(self, n_objectives: int, n_vectors: int) -> np.ndarray:
        """Generate uniformly distributed weight vectors"""
        if n_objectives == 2:
            # For 2 objectives, use simple linear distribution
            weights = np.zeros((n_vectors, 2))
            for i in range(n_vectors):
                w1 = i / (n_vectors - 1)
                weights[i] = [w1, 1 - w1]
            return weights
        else:
            # For higher dimensions, use random weights that sum to 1
            weights = np.random.random((n_vectors, n_objectives))
            weights = weights / np.sum(weights, axis=1, keepdims=True)
            return weights
    
    def _decompose_objectives(self, objective_values: np.ndarray,
                            weight_vectors: np.ndarray) -> np.ndarray:
        """Decompose multi-objective problem using weighted sum"""
        # Normalize objectives
        normalized_objectives = self._normalize_objectives(objective_values)
        
        # Calculate weighted sum for each weight vector and each solution
        decomposed_scores = np.dot(weight_vectors, normalized_objectives.T)
        
        return decomposed_scores
    
    def _normalize_objectives(self, objective_values: np.ndarray) -> np.ndarray:
        """Normalize objectives to [0, 1] range"""
        min_vals = np.min(objective_values, axis=0)
        max_vals = np.max(objective_values, axis=0)
        
        # Avoid division by zero
        ranges = max_vals - min_vals
        ranges[ranges == 0] = 1
        
        normalized = (objective_values - min_vals) / ranges
        return normalized
    
    def _evaluate_decomposition_quality(self, decomposed_scores: np.ndarray) -> float:
        """Evaluate quality of decomposition"""
        # Higher variance indicates better exploration of trade-offs
        return np.mean(np.var(decomposed_scores, axis=1))
    
    def _calculate_weight_diversity(self) -> float:
        """Calculate diversity of weight vectors"""
        if len(self.weight_vectors) < 2:
            return 0.0
        
        # Calculate pairwise distances between weight vectors
        distances = cdist(self.weight_vectors, self.weight_vectors)
        # Exclude diagonal (self-distances)
        distances = distances[np.triu_indices_from(distances, k=1)]
        
        return np.mean(distances)

class RandomForestMOOptimizer(BaseOptimizer):
    """
    Modern ML-based Multi-Objective Optimizer using Random Forest
    
    This represents the current state-of-the-art in ML-based compound optimization
    """
    
    def __init__(self, n_estimators: int = 200, max_depth: int = 20):
        super().__init__("RandomForest-MO")
        self.n_estimators = n_estimators
        self.max_depth = max_depth
        self.model = None
    
    def optimize(self, compounds_df: pd.DataFrame, objectives: Dict[str, Callable],
                max_iterations: int = 100) -> Tuple[pd.DataFrame, Dict]:
        """
        ML-based optimization using Random Forest
        """
        start_time = time.time()
        
        # Prepare features and targets
        feature_cols = ['molecular_weight', 'logp', 'hbd', 'hba', 'tpsa', 
                       'rotatable_bonds', 'aromatic_rings']
        
        available_features = [col for col in feature_cols if col in compounds_df.columns]
        
        if len(available_features) < 3:
            # Fallback to basic scoring
            return self._fallback_scoring(compounds_df, objectives)
        
        X = compounds_df[available_features].fillna(compounds_df[available_features].mean())
        
        # Calculate target objectives
        y_objectives = []
        objective_names = list(objectives.keys())
        
        for idx, row in compounds_df.iterrows():
            compound_dict = row.to_dict()
            obj_values = []
            
            for obj_func in objectives.values():
                try:
                    value = obj_func(compound_dict)
                    obj_values.append(value if isinstance(value, (int, float)) else 0.0)
                except:
                    obj_values.append(0.0)
            
            y_objectives.append(obj_values)
        
        y_objectives = np.array(y_objectives)
        
        # Train multi-output Random Forest
        self.model = MultiOutputRegressor(
            RandomForestRegressor(
                n_estimators=self.n_estimators,
                max_depth=self.max_depth,
                random_state=42,
                n_jobs=-1
            )
        )
        
        self.model.fit(X, y_objectives)
        
        # Predict objectives
        predicted_objectives = self.model.predict(X)
        
        # Calculate feature importance
        feature_importance = np.mean([
            estimator.feature_importances_ 
            for estimator in self.model.estimators_
        ], axis=0)
        
        # Multi-objective scoring using predicted values
        # Use Pareto efficiency for final scoring
        pareto_efficient = self._find_pareto_efficient(predicted_objectives)
        
        # Assign scores based on Pareto efficiency and prediction confidence
        scores = self._calculate_ml_scores(predicted_objectives, pareto_efficient, X)
        
        # Create scored dataframe
        scored_df = compounds_df.copy()
        scored_df['rf_mo_score'] = scores
        scored_df['pareto_efficient'] = pareto_efficient
        
        # Add predicted objectives
        for i, obj_name in enumerate(objective_names):
            scored_df[f'predicted_{obj_name}'] = predicted_objectives[:, i]
        
        scored_df = scored_df.sort_values('rf_mo_score', ascending=False)
        
        execution_time = time.time() - start_time
        
        # Calculate model performance metrics
        cv_scores = []
        for i in range(len(objective_names)):
            cv_score = cross_val_score(
                RandomForestRegressor(n_estimators=100, random_state=42),
                X, y_objectives[:, i], cv=5, scoring='r2'
            )
            cv_scores.append(np.mean(cv_score))
        
        metrics = {
            'algorithm': 'RandomForest-MO',
            'execution_time': execution_time,
            'n_pareto_efficient': np.sum(pareto_efficient),
            'model_performance': {
                'cv_r2_scores': cv_scores,
                'mean_cv_r2': np.mean(cv_scores),
                'feature_importance': dict(zip(available_features, feature_importance))
            },
            'prediction_quality': self._evaluate_prediction_quality(y_objectives, predicted_objectives)
        }
        
        return scored_df, metrics
    
    def _find_pareto_efficient(self, objectives: np.ndarray) -> np.ndarray:
        """Find Pareto efficient solutions"""
        n_points = objectives.shape[0]
        is_efficient = np.ones(n_points, dtype=bool)
        
        for i in range(n_points):
            if is_efficient[i]:
                # Check if any other point dominates this point
                dominated = np.all(objectives >= objectives[i], axis=1) & \
                           np.any(objectives > objectives[i], axis=1)
                is_efficient[dominated] = False
        
        return is_efficient
    
    def _calculate_ml_scores(self, predicted_objectives: np.ndarray,
                           pareto_efficient: np.ndarray, X: np.ndarray) -> np.ndarray:
        """Calculate ML-based scores combining prediction and efficiency"""
        
        # Base score from predicted objectives (weighted sum)
        weights = np.ones(predicted_objectives.shape[1]) / predicted_objectives.shape[1]
        base_scores = np.dot(predicted_objectives, weights)
        
        # Pareto efficiency bonus
        pareto_bonus = pareto_efficient.astype(float) * 0.2
        
        # Prediction confidence (based on feature values)
        confidence = self._calculate_prediction_confidence(X)
        
        # Final score
        final_scores = base_scores + pareto_bonus + confidence * 0.1
        
        return final_scores
    
    def _calculate_prediction_confidence(self, X: np.ndarray) -> np.ndarray:
        """Calculate prediction confidence based on feature space density"""
        # Simple confidence based on distance to training data centroid
        centroid = np.mean(X, axis=0)
        distances = np.linalg.norm(X - centroid, axis=1)
        max_distance = np.max(distances)
        
        # Higher confidence for points closer to centroid
        confidence = 1.0 - (distances / max_distance)
        
        return confidence
    
    def _evaluate_prediction_quality(self, true_objectives: np.ndarray,
                                   predicted_objectives: np.ndarray) -> Dict:
        """Evaluate prediction quality"""
        mse = np.mean((true_objectives - predicted_objectives) ** 2, axis=0)
        mae = np.mean(np.abs(true_objectives - predicted_objectives), axis=0)
        
        return {
            'mse_per_objective': mse.tolist(),
            'mae_per_objective': mae.tolist(),
            'overall_mse': np.mean(mse),
            'overall_mae': np.mean(mae)
        }
    
    def _fallback_scoring(self, compounds_df: pd.DataFrame, 
                         objectives: Dict[str, Callable]) -> Tuple[pd.DataFrame, Dict]:
        """Fallback scoring when insufficient features"""
        
        scores = []
        for idx, row in compounds_df.iterrows():
            compound_dict = row.to_dict()
            total_score = 0.0
            
            for obj_func in objectives.values():
                try:
                    score = obj_func(compound_dict)
                    total_score += score if isinstance(score, (int, float)) else 0.0
                except:
                    pass
            
            scores.append(total_score / len(objectives))
        
        scored_df = compounds_df.copy()
        scored_df['rf_mo_score'] = scores
        scored_df['pareto_efficient'] = False
        scored_df = scored_df.sort_values('rf_mo_score', ascending=False)
        
        metrics = {
            'algorithm': 'RandomForest-MO (Fallback)',
            'execution_time': 0.01,
            'note': 'Insufficient features for ML model'
        }
        
        return scored_df, metrics

class XGBoostMOOptimizer(BaseOptimizer):
    """
    XGBoost-based Multi-Objective Optimizer
    
    State-of-the-art gradient boosting for compound optimization
    """
    
    def __init__(self, n_estimators: int = 200, max_depth: int = 6, learning_rate: float = 0.1):
        super().__init__("XGBoost-MO")
        self.n_estimators = n_estimators
        self.max_depth = max_depth
        self.learning_rate = learning_rate
        self.models = {}
    
    def optimize(self, compounds_df: pd.DataFrame, objectives: Dict[str, Callable],
                max_iterations: int = 100) -> Tuple[pd.DataFrame, Dict]:
        """XGBoost-based optimization"""
        start_time = time.time()
        
        # Prepare features
        feature_cols = ['molecular_weight', 'logp', 'hbd', 'hba', 'tpsa', 
                       'rotatable_bonds', 'aromatic_rings']
        
        available_features = [col for col in feature_cols if col in compounds_df.columns]
        
        if len(available_features) < 3:
            return self._fallback_scoring(compounds_df, objectives)
        
        X = compounds_df[available_features].fillna(compounds_df[available_features].mean())
        
        # Train separate model for each objective
        objective_names = list(objectives.keys())
        predicted_objectives = np.zeros((len(compounds_df), len(objectives)))
        
        for i, (obj_name, obj_func) in enumerate(objectives.items()):
            # Calculate true objective values
            y_true = []
            for idx, row in compounds_df.iterrows():
                try:
                    value = obj_func(row.to_dict())
                    y_true.append(value if isinstance(value, (int, float)) else 0.0)
                except:
                    y_true.append(0.0)
            
            y_true = np.array(y_true)
            
            # Train XGBoost model
            model = xgb.XGBRegressor(
                n_estimators=self.n_estimators,
                max_depth=self.max_depth,
                learning_rate=self.learning_rate,
                random_state=42,
                n_jobs=-1
            )
            
            model.fit(X, y_true)
            self.models[obj_name] = model
            
            # Predict
            predicted_objectives[:, i] = model.predict(X)
        
        # Multi-objective scoring
        pareto_efficient = self._find_pareto_efficient(predicted_objectives)
        scores = self._calculate_xgb_scores(predicted_objectives, pareto_efficient)
        
        # Create scored dataframe
        scored_df = compounds_df.copy()
        scored_df['xgb_mo_score'] = scores
        scored_df['pareto_efficient'] = pareto_efficient
        
        for i, obj_name in enumerate(objective_names):
            scored_df[f'predicted_{obj_name}'] = predicted_objectives[:, i]
        
        scored_df = scored_df.sort_values('xgb_mo_score', ascending=False)
        
        execution_time = time.time() - start_time
        
        # Feature importance
        feature_importance = {}
        for obj_name, model in self.models.items():
            importance = model.feature_importances_
            feature_importance[obj_name] = dict(zip(available_features, importance))
        
        metrics = {
            'algorithm': 'XGBoost-MO',
            'execution_time': execution_time,
            'n_pareto_efficient': np.sum(pareto_efficient),
            'feature_importance': feature_importance,
            'model_performance': self._evaluate_xgb_performance(X, predicted_objectives, objectives, compounds_df)
        }
        
        return scored_df, metrics
    
    def _find_pareto_efficient(self, objectives: np.ndarray) -> np.ndarray:
        """Find Pareto efficient solutions (same as RandomForest)"""
        n_points = objectives.shape[0]
        is_efficient = np.ones(n_points, dtype=bool)
        
        for i in range(n_points):
            if is_efficient[i]:
                dominated = np.all(objectives >= objectives[i], axis=1) & \
                           np.any(objectives > objectives[i], axis=1)
                is_efficient[dominated] = False
        
        return is_efficient
    
    def _calculate_xgb_scores(self, predicted_objectives: np.ndarray,
                            pareto_efficient: np.ndarray) -> np.ndarray:
        """Calculate XGBoost-based scores"""
        # Weighted sum of predicted objectives
        weights = np.ones(predicted_objectives.shape[1]) / predicted_objectives.shape[1]
        base_scores = np.dot(predicted_objectives, weights)
        
        # Pareto efficiency bonus
        pareto_bonus = pareto_efficient.astype(float) * 0.3
        
        return base_scores + pareto_bonus
    
    def _evaluate_xgb_performance(self, X: np.ndarray, predicted_objectives: np.ndarray,
                                 objectives: Dict[str, Callable], compounds_df: pd.DataFrame) -> Dict:
        """Evaluate XGBoost model performance"""
        performance = {}
        
        for i, (obj_name, obj_func) in enumerate(objectives.items()):
            # Calculate true values
            y_true = []
            for idx, row in compounds_df.iterrows():
                try:
                    value = obj_func(row.to_dict())
                    y_true.append(value if isinstance(value, (int, float)) else 0.0)
                except:
                    y_true.append(0.0)
            
            y_true = np.array(y_true)
            y_pred = predicted_objectives[:, i]
            
            # Calculate metrics
            mse = np.mean((y_true - y_pred) ** 2)
            mae = np.mean(np.abs(y_true - y_pred))
            
            # R-squared
            ss_res = np.sum((y_true - y_pred) ** 2)
            ss_tot = np.sum((y_true - np.mean(y_true)) ** 2)
            r2 = 1 - (ss_res / ss_tot) if ss_tot != 0 else 0
            
            performance[obj_name] = {
                'mse': mse,
                'mae': mae,
                'r2': r2
            }
        
        return performance
    
    def _fallback_scoring(self, compounds_df: pd.DataFrame, 
                         objectives: Dict[str, Callable]) -> Tuple[pd.DataFrame, Dict]:
        """Fallback when insufficient features"""
        scores = []
        for idx, row in compounds_df.iterrows():
            total_score = 0.0
            for obj_func in objectives.values():
                try:
                    score = obj_func(row.to_dict())
                    total_score += score if isinstance(score, (int, float)) else 0.0
                except:
                    pass
            scores.append(total_score / len(objectives))
        
        scored_df = compounds_df.copy()
        scored_df['xgb_mo_score'] = scores
        scored_df = scored_df.sort_values('xgb_mo_score', ascending=False)
        
        return scored_df, {'algorithm': 'XGBoost-MO (Fallback)', 'execution_time': 0.01}

def get_all_baseline_optimizers() -> Dict[str, BaseOptimizer]:
    """Get all implemented baseline optimizers"""
    return {
        'nsga2': NSGAIIOptimizer(population_size=100),
        'moead': MOEADOptimizer(n_neighbors=10),
        'random_forest': RandomForestMOOptimizer(n_estimators=200),
        'xgboost': XGBoostMOOptimizer(n_estimators=200),
    }

if __name__ == "__main__":
    # Test the optimizers
    import pandas as pd
    
    # Create test data
    test_data = pd.DataFrame({
        'compound_id': [f'TEST_{i}' for i in range(100)],
        'molecular_weight': np.random.uniform(200, 600, 100),
        'logp': np.random.uniform(-1, 5, 100),
        'hbd': np.random.randint(0, 8, 100),
        'hba': np.random.randint(1, 12, 100),
        'tpsa': np.random.uniform(20, 150, 100),
        'rotatable_bonds': np.random.randint(0, 15, 100),
        'aromatic_rings': np.random.randint(0, 5, 100),
    })
    
    # Define test objectives
    def test_obj1(compound):
        return compound.get('molecular_weight', 0) / 500.0
    
    def test_obj2(compound):
        return 1.0 - abs(compound.get('logp', 0) - 2.5) / 5.0
    
    objectives = {'obj1': test_obj1, 'obj2': test_obj2}
    
    # Test all optimizers
    optimizers = get_all_baseline_optimizers()
    
    for name, optimizer in optimizers.items():
        print(f"\nTesting {name}...")
        scored_df, metrics = optimizer.optimize(test_data, objectives)
        print(f"  Score range: {scored_df.iloc[:, -1].min():.3f} - {scored_df.iloc[:, -1].max():.3f}")
        print(f"  Execution time: {metrics['execution_time']:.3f}s")
```
