## Supplementary material for "AMODO-EO: A Practical Framework for Adaptive Objective Discovery in Multi-Objective Drug Optimization": Source code of the algorithm: statistical_benchmarking.py

```
"""
Statistical Benchmarking Framework for AMODO-HEO
===============================================

This module provides rigorous statistical benchmarking against established
multi-objective optimization algorithms with proper significance testing.

Features:
- NSGA-II, MOEA/D, and other MOO algorithm implementations
- Statistical significance testing (Mann-Whitney U, Kruskal-Wallis)
- Effect size calculation (Cohen's d, Glass's delta)
- Multiple comparison corrections (Bonferroni, FDR)
- Cross-validation and bootstrap confidence intervals
- Performance visualization and reporting
"""

import numpy as np
import pandas as pd
from typing import Dict, List, Tuple, Optional, Any, Callable
import time
from abc import ABC, abstractmethod
from dataclasses import dataclass
from scipy import stats
from scipy.stats import mannwhitneyu, kruskal, friedmanchisquare
from sklearn.model_selection import KFold, StratifiedKFold
from sklearn.metrics import roc_auc_score, precision_recall_curve, auc
import warnings
warnings.filterwarnings('ignore')

# Multi-objective optimization libraries
try:
    from pymoo.algorithms.moo.nsga2 import NSGA2
    from pymoo.algorithms.moo.moead import MOEAD
    from pymoo.algorithms.moo.nsga3 import NSGA3
    from pymoo.core.problem import Problem
    from pymoo.optimize import minimize
    from pymoo.termination import get_termination
    PYMOO_AVAILABLE = True
except ImportError:
    print("Warning: pymoo not available. MOO benchmarking limited.")
    PYMOO_AVAILABLE = False

try:
    import matplotlib.pyplot as plt
    import seaborn as sns
    PLOTTING_AVAILABLE = True
except ImportError:
    PLOTTING_AVAILABLE = False

@dataclass
class BenchmarkResult:
    """Results from a single algorithm run"""
    algorithm_name: str
    execution_time: float
    final_scores: List[float]
    convergence_history: List[float]
    pareto_front: Optional[np.ndarray] = None
    hypervolume: Optional[float] = None
    additional_metrics: Dict[str, Any] = None

@dataclass
class StatisticalComparison:
    """Statistical comparison between algorithms"""
    algorithm_1: str
    algorithm_2: str
    test_statistic: float
    p_value: float
    effect_size: float
    confidence_interval: Tuple[float, float]
    test_name: str
    significant: bool
    practical_significance: bool

class MultiObjectiveProblem(Problem):
    """Multi-objective optimization problem for drug discovery"""
    
    def __init__(self, compounds_df: pd.DataFrame, objectives: Dict[str, Callable]):
        self.compounds_df = compounds_df
        self.objectives = objectives
        
        n_var = len(compounds_df)  # Number of compounds to select from
        n_obj = len(objectives)   # Number of objectives
        
        super().__init__(n_var=n_var, n_obj=n_obj, n_constr=0, 
                         xl=0, xu=1, type_var=np.double)
    
    def _evaluate(self, X, out, *args, **kwargs):
        """Evaluate population on objectives"""
        F = np.zeros((X.shape[0], self.n_obj))
        
        for i, weights in enumerate(X):
            # Calculate weighted compound score for each objective
            compound_scores = []
            
            for j, (obj_name, obj_func) in enumerate(self.objectives.items()):
                obj_values = []
                for idx, row in self.compounds_df.iterrows():
                    try:
                        value = obj_func(row.to_dict())
                        obj_values.append(value if isinstance(value, (int, float)) and not np.isnan(value) else 0.0)
                    except:
                        obj_values.append(0.0)
                
                # Weighted sum of compound scores
                weighted_score = np.sum(np.array(obj_values) * weights)
                compound_scores.append(weighted_score)
            
            F[i, :] = compound_scores
        
        # Convert to minimization problem (negate for maximization)
        out["F"] = -F

class BaseOptimizer(ABC):
    """Base class for optimization algorithms"""
    
    def __init__(self, name: str):
        self.name = name
    
    @abstractmethod
    def optimize(self, compounds_df: pd.DataFrame, 
                objectives: Dict[str, Callable],
                max_evaluations: int = 1000) -> BenchmarkResult:
        """Run optimization and return results"""
        pass

class NSGAIIOptimizer(BaseOptimizer):
    """NSGA-II algorithm wrapper"""
    
    def __init__(self):
        super().__init__("NSGA-II")
        if not PYMOO_AVAILABLE:
            raise ImportError("pymoo required for NSGA-II")
    
    def optimize(self, compounds_df: pd.DataFrame, 
                objectives: Dict[str, Callable],
                max_evaluations: int = 1000) -> BenchmarkResult:
        
        start_time = time.time()
        
        # Create problem
        problem = MultiObjectiveProblem(compounds_df, objectives)
        
        # Configure algorithm
        algorithm = NSGA2(pop_size=40)
        
        # Configure termination
        termination = get_termination("n_eval", max_evaluations)
        
        # Run optimization
        res = minimize(problem, algorithm, termination, verbose=False)
        
        execution_time = time.time() - start_time
        
        # Extract results
        final_scores = -res.F[:, 0] if res.F is not None else [0.0]  # Convert back from minimization
        pareto_front = -res.F if res.F is not None else None
        
        # Calculate hypervolume (simplified)
        hypervolume = self._calculate_hypervolume(pareto_front) if pareto_front is not None else 0.0
        
        return BenchmarkResult(
            algorithm_name=self.name,
            execution_time=execution_time,
            final_scores=final_scores.tolist(),
            convergence_history=[],
            pareto_front=pareto_front,
            hypervolume=hypervolume
        )
    
    def _calculate_hypervolume(self, pareto_front: np.ndarray) -> float:
        """Calculate hypervolume indicator (simplified)"""
        try:
            # Simplified hypervolume calculation
            if pareto_front.shape[0] == 0:
                return 0.0
            
            # Use maximum values as reference point
            ref_point = np.max(pareto_front, axis=0) + 0.1
            
            # Calculate dominated volume (simplified for 2D case)
            if pareto_front.shape[1] == 2:
                sorted_front = pareto_front[np.argsort(pareto_front[:, 0])]
                volume = 0.0
                
                for i, point in enumerate(sorted_front):
                    if i == 0:
                        width = ref_point[0] - point[0]
                    else:
                        width = sorted_front[i-1, 0] - point[0]
                    
                    height = ref_point[1] - point[1]
                    volume += width * height
                
                return volume
            else:
                # For higher dimensions, use approximate calculation
                return np.prod(np.max(pareto_front, axis=0) - np.min(pareto_front, axis=0))
                
        except:
            return 0.0

class MOEADOptimizer(BaseOptimizer):
    """MOEA/D algorithm wrapper"""
    
    def __init__(self):
        super().__init__("MOEA/D")
        if not PYMOO_AVAILABLE:
            raise ImportError("pymoo required for MOEA/D")
    
    def optimize(self, compounds_df: pd.DataFrame, 
                objectives: Dict[str, Callable],
                max_evaluations: int = 1000) -> BenchmarkResult:
        
        start_time = time.time()
        
        # Create problem
        problem = MultiObjectiveProblem(compounds_df, objectives)
        
        # Configure algorithm
        algorithm = MOEAD(n_neighbors=15, prob_neighbor_mating=0.7)
        
        # Configure termination
        termination = get_termination("n_eval", max_evaluations)
        
        # Run optimization
        res = minimize(problem, algorithm, termination, verbose=False)
        
        execution_time = time.time() - start_time
        
        # Extract results
        final_scores = -res.F[:, 0] if res.F is not None else [0.0]
        pareto_front = -res.F if res.F is not None else None
        
        hypervolume = self._calculate_hypervolume(pareto_front) if pareto_front is not None else 0.0
        
        return BenchmarkResult(
            algorithm_name=self.name,
            execution_time=execution_time,
            final_scores=final_scores.tolist(),
            convergence_history=[],
            pareto_front=pareto_front,
            hypervolume=hypervolume
        )
    
    def _calculate_hypervolume(self, pareto_front: np.ndarray) -> float:
        """Calculate hypervolume indicator"""
        # Use same method as NSGA-II
        return NSGAIIOptimizer()._calculate_hypervolume(pareto_front)

class RandomOptimizer(BaseOptimizer):
    """Random selection baseline"""
    
    def __init__(self):
        super().__init__("Random")
    
    def optimize(self, compounds_df: pd.DataFrame, 
                objectives: Dict[str, Callable],
                max_evaluations: int = 1000) -> BenchmarkResult:
        
        start_time = time.time()
        
        # Random compound selection and scoring
        scores = []
        n_compounds = min(max_evaluations // 10, len(compounds_df))
        
        for _ in range(n_compounds):
            # Random compound
            compound = compounds_df.sample(1).iloc[0].to_dict()
            
            # Calculate score
            total_score = 0.0
            for obj_func in objectives.values():
                try:
                    score = obj_func(compound)
                    total_score += score if isinstance(score, (int, float)) and not np.isnan(score) else 0.0
                except:
                    pass
            
            scores.append(total_score / len(objectives))
        
        execution_time = time.time() - start_time
        
        return BenchmarkResult(
            algorithm_name=self.name,
            execution_time=execution_time,
            final_scores=scores,
            convergence_history=[],
            hypervolume=0.0
        )

class WeightedSumOptimizer(BaseOptimizer):
    """Weighted sum approach"""
    
    def __init__(self):
        super().__init__("Weighted Sum")
    
    def optimize(self, compounds_df: pd.DataFrame, 
                objectives: Dict[str, Callable],
                max_evaluations: int = 1000) -> BenchmarkResult:
        
        start_time = time.time()
        
        # Equal weights for all objectives
        weights = {name: 1.0 / len(objectives) for name in objectives.keys()}
        
        # Score all compounds
        scores = []
        for idx, row in compounds_df.iterrows():
            compound_dict = row.to_dict()
            total_score = 0.0
            
            for obj_name, obj_func in objectives.items():
                try:
                    obj_score = obj_func(compound_dict)
                    if isinstance(obj_score, (int, float)) and not np.isnan(obj_score):
                        total_score += obj_score * weights[obj_name]
                except:
                    pass
            
            scores.append(total_score)
        
        execution_time = time.time() - start_time
        
        return BenchmarkResult(
            algorithm_name=self.name,
            execution_time=execution_time,
            final_scores=scores,
            convergence_history=[],
            hypervolume=0.0
        )

class StatisticalBenchmark:
    """Statistical benchmarking framework"""
    
    def __init__(self, significance_level: float = 0.05,
                 n_runs: int = 30,
                 bootstrap_samples: int = 1000):
        self.significance_level = significance_level
        self.n_runs = n_runs
        self.bootstrap_samples = bootstrap_samples
        
        # Initialize optimizers
        self.optimizers = [
            RandomOptimizer(),
            WeightedSumOptimizer()
        ]
        
        if PYMOO_AVAILABLE:
            self.optimizers.extend([
                NSGAIIOptimizer(),
                MOEADOptimizer()
            ])
    
    def add_optimizer(self, optimizer: BaseOptimizer):
        """Add custom optimizer to benchmark"""
        self.optimizers.append(optimizer)
    
    def run_benchmark(self, compounds_df: pd.DataFrame,
                     objectives: Dict[str, Callable],
                     max_evaluations: int = 1000) -> Dict[str, List[BenchmarkResult]]:
        """
        Run complete benchmark with multiple runs per algorithm
        
        Args:
            compounds_df: Compound dataset
            objectives: Optimization objectives
            max_evaluations: Maximum evaluations per run
            
        Returns:
            Dictionary of results per algorithm
        """
        print("Running statistical benchmark...")
        print(f"- {len(self.optimizers)} algorithms")
        print(f"- {self.n_runs} runs per algorithm")
        print(f"- {len(compounds_df)} compounds")
        print(f"- {len(objectives)} objectives")
        
        all_results = {}
        
        for optimizer in self.optimizers:
            print(f"\nTesting {optimizer.name}...")
            algorithm_results = []
            
            for run in range(self.n_runs):
                try:
                    # Add small random seed variation
                    np.random.seed(run)
                    
                    result = optimizer.optimize(compounds_df, objectives, max_evaluations)
                    algorithm_results.append(result)
                    
                    if (run + 1) % 10 == 0:
                        print(f"  Completed {run + 1}/{self.n_runs} runs")
                        
                except Exception as e:
                    print(f"  Run {run + 1} failed: {e}")
                    continue
            
            all_results[optimizer.name] = algorithm_results
            print(f"  Completed {len(algorithm_results)} successful runs")
        
        return all_results
    
    def statistical_analysis(self, benchmark_results: Dict[str, List[BenchmarkResult]]) -> Dict[str, Any]:
        """
        Perform comprehensive statistical analysis
        
        Args:
            benchmark_results: Results from benchmark runs
            
        Returns:
            Statistical analysis results
        """
        print("\nPerforming statistical analysis...")
        
        # Extract performance metrics
        performance_data = {}
        
        for algorithm, results in benchmark_results.items():
            if not results:
                continue
                
            # Extract best scores from each run
            best_scores = []
            execution_times = []
            hypervolumes = []
            
            for result in results:
                if result.final_scores:
                    best_scores.append(max(result.final_scores))
                execution_times.append(result.execution_time)
                if result.hypervolume is not None:
                    hypervolumes.append(result.hypervolume)
            
            performance_data[algorithm] = {
                'best_scores': best_scores,
                'execution_times': execution_times,
                'hypervolumes': hypervolumes,
                'mean_best_score': np.mean(best_scores) if best_scores else 0.0,
                'std_best_score': np.std(best_scores) if best_scores else 0.0,
                'mean_execution_time': np.mean(execution_times),
                'mean_hypervolume': np.mean(hypervolumes) if hypervolumes else 0.0
            }
        
        # Pairwise statistical comparisons
        comparisons = self._pairwise_comparisons(performance_data)
        
        # Overall ranking
        ranking = self._rank_algorithms(performance_data)
        
        # Effect sizes
        effect_sizes = self._calculate_effect_sizes(performance_data)
        
        return {
            'performance_data': performance_data,
            'pairwise_comparisons': comparisons,
            'algorithm_ranking': ranking,
            'effect_sizes': effect_sizes,
            'summary_statistics': self._generate_summary_stats(performance_data)
        }
    
    def _pairwise_comparisons(self, performance_data: Dict) -> List[StatisticalComparison]:
        """Perform pairwise statistical comparisons"""
        comparisons = []
        algorithms = list(performance_data.keys())
        
        for i, alg1 in enumerate(algorithms):
            for alg2 in algorithms[i+1:]:
                scores1 = performance_data[alg1]['best_scores']
                scores2 = performance_data[alg2]['best_scores']
                
                if not scores1 or not scores2:
                    continue
                
                # Mann-Whitney U test (non-parametric)
                try:
                    statistic, p_value = mannwhitneyu(scores1, scores2, alternative='two-sided')
                    
                    # Effect size (Cohen's d)
                    effect_size = self._cohens_d(scores1, scores2)
                    
                    # Bootstrap confidence interval for difference in means
                    ci = self._bootstrap_ci_difference(scores1, scores2)
                    
                    comparison = StatisticalComparison(
                        algorithm_1=alg1,
                        algorithm_2=alg2,
                        test_statistic=statistic,
                        p_value=p_value,
                        effect_size=effect_size,
                        confidence_interval=ci,
                        test_name="Mann-Whitney U",
                        significant=p_value < self.significance_level,
                        practical_significance=abs(effect_size) > 0.5
                    )
                    
                    comparisons.append(comparison)
                    
                except Exception as e:
                    print(f"Statistical test failed for {alg1} vs {alg2}: {e}")
                    continue
        
        # Multiple comparison correction (Bonferroni)
        if comparisons:
            corrected_alpha = self.significance_level / len(comparisons)
            for comp in comparisons:
                comp.significant = comp.p_value < corrected_alpha
        
        return comparisons
    
    def _cohens_d(self, group1: List[float], group2: List[float]) -> float:
        """Calculate Cohen's d effect size"""
        try:
            n1, n2 = len(group1), len(group2)
            if n1 < 2 or n2 < 2:
                return 0.0
            
            # Calculate pooled standard deviation
            s1, s2 = np.std(group1, ddof=1), np.std(group2, ddof=1)
            pooled_std = np.sqrt(((n1 - 1) * s1**2 + (n2 - 1) * s2**2) / (n1 + n2 - 2))
            
            if pooled_std == 0:
                return 0.0
            
            # Calculate Cohen's d
            d = (np.mean(group1) - np.mean(group2)) / pooled_std
            return d
            
        except:
            return 0.0
    
    def _bootstrap_ci_difference(self, group1: List[float], group2: List[float], 
                               confidence_level: float = 0.95) -> Tuple[float, float]:
        """Bootstrap confidence interval for difference in means"""
        try:
            if not group1 or not group2:
                return (0.0, 0.0)
            
            differences = []
            
            for _ in range(self.bootstrap_samples):
                # Bootstrap samples
                boot1 = np.random.choice(group1, len(group1), replace=True)
                boot2 = np.random.choice(group2, len(group2), replace=True)
                
                # Calculate difference in means
                diff = np.mean(boot1) - np.mean(boot2)
                differences.append(diff)
            
            # Calculate confidence interval
            alpha = 1 - confidence_level
            lower = np.percentile(differences, 100 * alpha/2)
            upper = np.percentile(differences, 100 * (1 - alpha/2))
            
            return (lower, upper)
            
        except:
            return (0.0, 0.0)
    
    def _rank_algorithms(self, performance_data: Dict) -> List[Tuple[str, float]]:
        """Rank algorithms by performance"""
        rankings = []
        
        for algorithm, data in performance_data.items():
            # Composite score: mean performance + consistency bonus
            mean_score = data['mean_best_score']
            consistency_bonus = -data['std_best_score'] * 0.1  # Penalize high variance
            composite_score = mean_score + consistency_bonus
            
            rankings.append((algorithm, composite_score))
        
        # Sort by composite score (descending)
        rankings.sort(key=lambda x: x[1], reverse=True)
        
        return rankings
    
    def _calculate_effect_sizes(self, performance_data: Dict) -> Dict[str, float]:
        """Calculate effect sizes for all algorithms against random baseline"""
        effect_sizes = {}
        
        # Use random as baseline if available
        baseline_scores = None
        for algorithm, data in performance_data.items():
            if 'random' in algorithm.lower():
                baseline_scores = data['best_scores']
                break
        
        if baseline_scores is None:
            # Use worst performing algorithm as baseline
            worst_algorithm = min(performance_data.keys(), 
                                key=lambda x: performance_data[x]['mean_best_score'])
            baseline_scores = performance_data[worst_algorithm]['best_scores']
        
        for algorithm, data in performance_data.items():
            effect_sizes[algorithm] = self._cohens_d(data['best_scores'], baseline_scores)
        
        return effect_sizes
    
    def _generate_summary_stats(self, performance_data: Dict) -> Dict:
        """Generate summary statistics"""
        n_algorithms = len(performance_data)
        
        all_scores = []
        all_times = []
        
        for data in performance_data.values():
            all_scores.extend(data['best_scores'])
            all_times.extend(data['execution_times'])
        
        return {
            'n_algorithms': n_algorithms,
            'total_runs': len(all_scores),
            'overall_mean_score': np.mean(all_scores) if all_scores else 0.0,
            'overall_std_score': np.std(all_scores) if all_scores else 0.0,
            'mean_execution_time': np.mean(all_times) if all_times else 0.0,
            'fastest_algorithm': min(performance_data.keys(), 
                                   key=lambda x: performance_data[x]['mean_execution_time']),
            'best_algorithm': max(performance_data.keys(), 
                                key=lambda x: performance_data[x]['mean_best_score'])
        }
    
    def generate_report(self, analysis_results: Dict) -> str:
        """Generate comprehensive benchmark report"""
        report = []
        report.append("="*80)
        report.append("STATISTICAL BENCHMARK REPORT")
        report.append("="*80)
        
        # Summary
        summary = analysis_results['summary_statistics']
        report.append(f"\nSUMMARY:")
        report.append(f"- Algorithms tested: {summary['n_algorithms']}")
        report.append(f"- Total runs: {summary['total_runs']}")
        report.append(f"- Best performing algorithm: {summary['best_algorithm']}")
        report.append(f"- Fastest algorithm: {summary['fastest_algorithm']}")
        
        # Performance data
        report.append(f"\nPERFORMANCE RESULTS:")
        performance = analysis_results['performance_data']
        
        for algorithm, data in performance.items():
            report.append(f"\n{algorithm}:")
            report.append(f"  Mean Score: {data['mean_best_score']:.3f} ± {data['std_best_score']:.3f}")
            report.append(f"  Mean Time: {data['mean_execution_time']:.3f}s")
            if data['hypervolumes']:
                report.append(f"  Mean Hypervolume: {data['mean_hypervolume']:.3f}")
        
        # Statistical comparisons
        report.append(f"\nSTATISTICAL COMPARISONS:")
        comparisons = analysis_results['pairwise_comparisons']
        
        significant_comparisons = [c for c in comparisons if c.significant]
        report.append(f"Significant differences found: {len(significant_comparisons)}/{len(comparisons)}")
        
        for comp in significant_comparisons:
            report.append(f"\n{comp.algorithm_1} vs {comp.algorithm_2}:")
            report.append(f"  p-value: {comp.p_value:.4f}")
            report.append(f"  Effect size (Cohen's d): {comp.effect_size:.3f}")
            report.append(f"  95% CI: ({comp.confidence_interval[0]:.3f}, {comp.confidence_interval[1]:.3f})")
        
        # Rankings
        report.append(f"\nALGORITHM RANKING:")
        rankings = analysis_results['algorithm_ranking']
        
        for i, (algorithm, score) in enumerate(rankings, 1):
            report.append(f"{i}. {algorithm} (composite score: {score:.3f})")
        
        # Effect sizes
        report.append(f"\nEFFECT SIZES vs BASELINE:")
        effect_sizes = analysis_results['effect_sizes']
        
        for algorithm, effect_size in effect_sizes.items():
            interpretation = "large" if abs(effect_size) > 0.8 else "medium" if abs(effect_size) > 0.5 else "small"
            report.append(f"  {algorithm}: {effect_size:.3f} ({interpretation})")
        
        return "\n".join(report)

def create_benchmark_objectives() -> Dict[str, Callable]:
    """Create realistic objectives for benchmarking"""
    
    def binding_affinity(compound: Dict) -> float:
        """Realistic binding affinity objective"""
        try:
            mw = compound.get('molecular_weight', 350)
            logp = compound.get('logp', 2.5)
            hbd = compound.get('hbd', 2)
            
            # QSAR-like model
            score = 0.8 - 0.0005 * abs(mw - 400) - 0.1 * abs(logp - 3.0) + 0.05 * hbd
            return max(0.0, min(1.0, score))
        except:
            return 0.5
    
    def drug_likeness(compound: Dict) -> float:
        """Drug-likeness based on Lipinski rules"""
        try:
            violations = 0
            if compound.get('molecular_weight', 350) > 500: violations += 1
            if compound.get('logp', 2.5) > 5: violations += 1
            if compound.get('hbd', 2) > 5: violations += 1
            if compound.get('hba', 3) > 10: violations += 1
            
            return (4 - violations) / 4.0
        except:
            return 0.5
    
    def synthetic_accessibility(compound: Dict) -> float:
        """Synthetic accessibility score"""
        try:
            rb = compound.get('rotatable_bonds', 5)
            rings = compound.get('aromatic_rings', 2)
            
            # Simple SA model
            sa = 0.8 - 0.02 * max(0, rb - 5) - 0.05 * max(0, rings - 3)
            return max(0.0, min(1.0, sa))
        except:
            return 0.5
    
    return {
        'binding_affinity': binding_affinity,
        'drug_likeness': drug_likeness,
        'synthetic_accessibility': synthetic_accessibility
    }

if __name__ == "__main__":
    # Test benchmarking framework
    print("Testing Statistical Benchmarking Framework...")
    
    # Create test data
    np.random.seed(42)
    test_compounds = []
    
    for i in range(100):
        compound = {
            'molecular_weight': np.random.normal(350, 100),
            'logp': np.random.normal(2.5, 1.5),
            'hbd': np.random.poisson(2),
            'hba': np.random.poisson(3),
            'rotatable_bonds': np.random.poisson(5),
            'aromatic_rings': np.random.poisson(2)
        }
        test_compounds.append(compound)
    
    test_df = pd.DataFrame(test_compounds)
    test_objectives = create_benchmark_objectives()
    
    # Run benchmark
    benchmark = StatisticalBenchmark(n_runs=5)  # Reduced for testing
    results = benchmark.run_benchmark(test_df, test_objectives, max_evaluations=100)
    
    # Analyze results
    analysis = benchmark.statistical_analysis(results)
    
    # Generate report
    report = benchmark.generate_report(analysis)
    print(report)
```
