## Supplementary material for "AMODO-EO: A Practical Framework for Adaptive Objective Discovery in Multi-Objective Drug Optimization": Source code of the algorithm: test_json_fix.py

```
"""
Test JSON serialization fix
"""

import sys
sys.path.append('src')

from src.comprehensive_benchmark import ComprehensiveBenchmark

print("Testing JSON serialization fix...")

# Initialize benchmark
benchmark = ComprehensiveBenchmark(results_dir="results/test_json_fix")

print("Running small benchmark...")

# Run minimal benchmark
try:
    results = benchmark.run_comprehensive_benchmark(
        n_folds=2,  # Reduced for quick test
        datasets=['benchmark'],  # Just one dataset  
        algorithms=['amodo_heo', 'random']  # Just two algorithms
    )
    print("✅ JSON serialization test passed!")
    print(f"Publication ready: {results.get('publication_ready', {}).get('publication_ready', 'Unknown')}")
    
except Exception as e:
    print(f"❌ Test failed: {e}")
    import traceback
    traceback.print_exc()
```
