## Supplementary material for "AMODO-EO: A Practical Framework for Adaptive Objective Discovery in Multi-Objective Drug Optimization": Source code of the algorithm: test_publication_readiness.py

```
"""
Quick Publication Readiness Test
===============================

This script runs a quick test to verify the publication-ready AMODO-HEO system
is working correctly before running the full benchmark.

Author: Nihal Shetty
Date: August 4, 2025
"""

import sys
import os
import traceback
import time
import json
import pandas as pd
from pathlib import Path

def test_imports():
    """Test if all required modules can be imported"""
    print("Testing imports...")
    
    try:
        # Standard libraries
        import numpy as np
        import pandas as pd
        import scipy
        from sklearn.model_selection import cross_val_score
        from sklearn.ensemble import RandomForestRegressor
        print("✅ Standard libraries imported successfully")
        
        # Try to import our custom modules
        sys.path.append('src')
        
        # Test dataset management
        try:
            from src.real_datasets import RealDatasetManager
            print("✅ RealDatasetManager imported successfully")
        except ImportError as e:
            print(f"❌ Failed to import RealDatasetManager: {e}")
            return False
        
        # Test baseline algorithms
        try:
            from src.modern_baselines import get_all_baseline_optimizers, NSGAIIOptimizer
            print("✅ Modern baselines imported successfully")
        except ImportError as e:
            print(f"❌ Failed to import modern baselines: {e}")
            return False
        
        # Test comprehensive benchmark
        try:
            from src.comprehensive_benchmark import ComprehensiveBenchmark
            print("✅ ComprehensiveBenchmark imported successfully")
        except ImportError as e:
            print(f"❌ Failed to import ComprehensiveBenchmark: {e}")
            return False
        
        # Test main algorithm
        try:
            from src.publication_ready_amodo_heo import PublicationReadyAMODOHEO
            print("✅ PublicationReadyAMODOHEO imported successfully")
        except ImportError as e:
            print(f"❌ Failed to import PublicationReadyAMODOHEO: {e}")
            return False
        
        return True
        
    except ImportError as e:
        print(f"❌ Import error: {e}")
        return False

def test_dataset_creation():
    """Test dataset creation"""
    print("\nTesting dataset creation...")
    
    try:
        from src.real_datasets import RealDatasetManager
        
        # Create small test dataset
        dataset_manager = RealDatasetManager()
        
        # Test benchmark dataset creation
        X, y = dataset_manager.get_dataset('benchmark', n_compounds=100)
        
        if X is not None and y is not None:
            print(f"✅ Created benchmark dataset: {X.shape[0]} compounds, {X.shape[1]} features, {y.shape[1]} objectives")
            return True
        else:
            print("❌ Failed to create benchmark dataset")
            return False
            
    except Exception as e:
        print(f"❌ Dataset creation failed: {e}")
        print(traceback.format_exc())
        return False

def test_baseline_algorithms():
    """Test baseline algorithm initialization"""
    print("\nTesting baseline algorithms...")
    
    try:
        from src.modern_baselines import get_all_baseline_optimizers
        
        # Get all optimizers
        optimizers = get_all_baseline_optimizers()
        
        print(f"✅ Created {len(optimizers)} baseline optimizers:")
        for name in optimizers.keys():
            print(f"    • {name}")
        
        return True
        
    except Exception as e:
        print(f"❌ Baseline algorithm test failed: {e}")
        print(traceback.format_exc())
        return False

def test_amodo_heo_algorithm():
    """Test AMODO-HEO algorithm initialization"""
    print("\nTesting AMODO-HEO algorithm...")
    
    try:
        from src.publication_ready_amodo_heo import PublicationReadyAMODOHEO
        
        # Initialize algorithm
        amodo = PublicationReadyAMODOHEO(
            max_objectives=3,
            significance_threshold=0.1,
            min_improvement=0.01
        )
        
        print(f"✅ AMODO-HEO initialized successfully")
        print(f"    Max objectives: {amodo.max_objectives}")
        print(f"    Significance threshold: {amodo.significance_threshold}")
        print(f"    Min improvement: {amodo.min_improvement}")
        
        return True
        
    except Exception as e:
        print(f"❌ AMODO-HEO test failed: {e}")
        print(traceback.format_exc())
        return False

def test_quick_optimization():
    """Test a quick optimization run"""
    print("\nTesting quick optimization...")
    
    try:
        from src.real_datasets import RealDatasetManager
        from src.publication_ready_amodo_heo import PublicationReadyAMODOHEO
        
        # Create small dataset
        dataset_manager = RealDatasetManager()
        X, y = dataset_manager.get_dataset('benchmark', n_compounds=50)
        
        # Initialize AMODO-HEO with minimal parameters
        amodo = PublicationReadyAMODOHEO(
            max_objectives=y.shape[1],
            significance_threshold=0.1,
            min_improvement=0.01
        )
        
        print("Running quick optimization...")
        start_time = time.time()
        
        # Run optimization (check if optimize method exists)
        if hasattr(amodo, 'optimize'):
            pareto_front = amodo.optimize(X, y)
        elif hasattr(amodo, 'optimize_compounds'):
            # Create a DataFrame for optimize_compounds method
            df = pd.DataFrame(X, columns=[f'feature_{i}' for i in range(X.shape[1])])
            objectives = [f'objective_{i}' for i in range(y.shape[1])]
            pareto_front, _ = amodo.optimize_compounds(df, objectives)
        else:
            print("❌ No optimization method found")
            return False
        
        elapsed_time = time.time() - start_time
        
        if pareto_front is not None and len(pareto_front) > 0:
            print(f"✅ Quick optimization completed in {elapsed_time:.2f} seconds")
            print(f"    Pareto front size: {len(pareto_front)}")
            return True
        else:
            print("❌ Optimization failed - empty Pareto front")
            return False
        
    except Exception as e:
        print(f"❌ Quick optimization failed: {e}")
        print(traceback.format_exc())
        return False

def test_benchmark_framework():
    """Test benchmark framework initialization"""
    print("\nTesting benchmark framework...")
    
    try:
        from src.comprehensive_benchmark import ComprehensiveBenchmark
        
        # Initialize benchmark
        benchmark = ComprehensiveBenchmark(results_dir="results/test_benchmark")
        
        print(f"✅ Benchmark framework initialized")
        print(f"    Results directory: {benchmark.results_dir}")
        
        return True
        
    except Exception as e:
        print(f"❌ Benchmark framework test failed: {e}")
        print(traceback.format_exc())
        return False

def run_all_tests():
    """Run all tests"""
    print("="*80)
    print("AMODO-HEO PUBLICATION READINESS - QUICK TEST")
    print("="*80)
    print("Version: 2.0")
    print("Date: August 4, 2025")
    print("="*80)
    
    tests = [
        ("Import Test", test_imports),
        ("Dataset Creation", test_dataset_creation),
        ("Baseline Algorithms", test_baseline_algorithms),
        ("AMODO-HEO Algorithm", test_amodo_heo_algorithm),
        ("Quick Optimization", test_quick_optimization),
        ("Benchmark Framework", test_benchmark_framework),
    ]
    
    results = {}
    total_tests = len(tests)
    passed_tests = 0
    
    for test_name, test_func in tests:
        print(f"\n{'='*20} {test_name} {'='*20}")
        
        try:
            result = test_func()
            results[test_name] = result
            if result:
                passed_tests += 1
                print(f"✅ {test_name}: PASSED")
            else:
                print(f"❌ {test_name}: FAILED")
        except Exception as e:
            print(f"❌ {test_name}: ERROR - {e}")
            results[test_name] = False
    
    # Final results
    print("\n" + "="*80)
    print("TEST RESULTS SUMMARY")
    print("="*80)
    
    success_rate = (passed_tests / total_tests) * 100
    
    print(f"Tests Passed: {passed_tests}/{total_tests} ({success_rate:.1f}%)")
    
    if success_rate >= 100:
        print("🎉 ALL TESTS PASSED! System is ready for publication benchmark!")
        print("\nNext steps:")
        print("  1. Run: python run_publication_benchmark.py")
        print("  2. This will generate publication-ready results")
        print("  3. Review results in results/publication_benchmark/")
        
    elif success_rate >= 80:
        print("⚠️ Most tests passed, but some issues detected")
        print("System should work but may have minor issues")
        
        print("\nFailed tests:")
        for test_name, result in results.items():
            if not result:
                print(f"  • {test_name}")
        
    else:
        print("❌ MAJOR ISSUES DETECTED")
        print("System is not ready for publication benchmark")
        
        print("\nRequired fixes:")
        for test_name, result in results.items():
            if not result:
                print(f"  • Fix {test_name}")
        
        print("\nTroubleshooting:")
        print("  1. Check that all dependencies are installed: pip install -r requirements.txt")
        print("  2. Ensure all src/ files are present")
        print("  3. Check file permissions and paths")
    
    print(f"\nOverall Status: {'✅ READY' if success_rate >= 100 else '⚠️ ISSUES' if success_rate >= 80 else '❌ NOT READY'}")
    
    return results

if __name__ == "__main__":
    results = run_all_tests()
    
    # Save test results
    os.makedirs("results", exist_ok=True)
    with open("results/test_results.json", "w") as f:
        json.dump({
            "timestamp": time.strftime("%Y%m%d_%H%M%S"),
            "test_results": results,
            "summary": {
                "total_tests": len(results),
                "passed_tests": sum(results.values()),
                "success_rate": (sum(results.values()) / len(results)) * 100
            }
        }, f, indent=2)
    
    print(f"\n📄 Test results saved to: results/test_results.json")
```
